# Inhibition of a cortico-thalamic circuit attenuates cue-induced reinstatement of drug-seeking behavior in “relapse prone” male rats

**DOI:** 10.1101/2020.03.01.972224

**Authors:** Brittany N. Kuhn, Paolo Campus, Marin S. Klumpner, Stephen E. Chang, Amanda G. Iglesias, Shelly B. Flagel

## Abstract

Relapse often occurs when individuals are exposed to stimuli or cues previously associated with the drug-taking experience. The ability of drug cues to trigger relapse is believed to be a consequence of incentive salience attribution, a process by which the incentive value of reward is transferred to the reward-paired cue. Sign-tracker (ST) rats that attribute enhanced incentive value to reward cues are more prone to relapse compared to goal-tracker (GT) rats that primarily attribute predictive value to such cues. The neurobiological mechanisms underlying this individual variation in relapse propensity remains largely unexplored. The paraventricular nucleus of the thalamus (PVT) has been identified as a critical node in the regulation of cue-elicited behaviors in STs and GTs, including cue-induced reinstatement of drug-seeking behavior. Here we used a chemogenetic approach to assess whether “top-down” cortical input from the prelimbic cortex (PrL) to the PVT plays a role in mediating individual differences in relapse propensity. Chemogenetic inhibition of the PrL-PVT pathway selectively decreased cue-induced reinstatement of drug-seeking behavior in STs, without affecting behavior in GTs. In contrast, cocaine-primed drug-seeking behavior was not affected in either phenotype. Furthermore, when rats were characterized based on a different behavioral phenotype – locomotor response to novelty – inhibition of the PrL-PVT pathway had no effect on either cue- or drug-induced reinstatement. These results highlight an important role for the PrL-PVT pathway in vulnerability to relapse that is driven by individual differences in the propensity to attribute incentive salience to discrete reward cues.

## Introduction

Relapse remains one of the biggest challenges in the treatment of drug addiction [1], with relapse rates as high as 75% within one year of treatment for cocaine addiction [2]. Individuals with addiction often relapse upon exposure to cues (e.g. people, places, paraphernalia) associated with their prior drug-taking experience. Such environmental stimuli can acquire the ability to elicit craving [3, 4] and instigate drug-seeking behavior [5] through Pavlovian learning processes. When a cue reliably precedes the delivery of a reward, it is attributed with predictive value; but Pavlovian cues can also acquire excessive motivational value, or incentive salience [6]. The extent to which a reward cue is attributed with incentive value varies between individuals. Only for some individuals are reward cues transformed into incentive stimuli or “motivational magnets” [7] and, when this occurs, such cues attain inordinate control and invigorate maladaptive behaviors, including drug seeking and drug taking.

Individual variation in cue-reward learning can be assessed using the sign-tracker/ goal-tracker animal model [8–12]. When rats are exposed to a Pavlovian conditioned approach procedure, wherein a discrete cue precedes the delivery of a food reward, different phenotypes emerge. Some rats, referred to as goal-trackers (GTs) [13], attribute only predictive value to the reward cue; others, termed sign-trackers (STs) [14], attribute both predictive and incentive value to the cue [8, 9, 12]. STs and GTs also differ on several addiction-related behaviors [15–25]. Relative to GTs, STs show greater drug-seeking behavior during tests for cocaine-primed [19] or cue-induced reinstatement [20, 21] following limited drug experience and abstinence. These phenotypic differences in relapse propensity support the long-standing belief that Pavlovian learning processes [6, 26–29], and, more explicitly, incentive salience attribution [6], contribute to addiction. The ST/GT model, therefore, provides a means to elucidate the neurobiological mechanisms underlying the incentive motivational processes that contribute to relapse propensity.

The behavioral phenotypes exhibited by sign-trackers and goal-trackers have been postulated to result from a neurobiological bias between “bottom-up” and “top-down” processes, respectively [23, 25, 30–33]. The paraventricular nucleus of the thalamus (PVT) may play a critical role in this regard, as it is ideally located to receive cortical and subcortical input and regulate motivated behaviors via output to the ventral striatum [34–37]. The PVT has long-been recognized as a node of the motive circuit [38, for review see 39], and more recently for its role in addiction-like behaviors, including drug seeking [40–43]. We have previously shown that inactivation of the PVT robustly increases cue-induced drug-seeking behavior in GTs, without affecting behavior in STs [44]. The PVT receives dense projections from the prelimbic (PrL) cortex, and both phenotypes engage this “top-down” pathway in response to a food cue, suggesting that it mediates the predictive value of a Pavlovian reward cue [32]. Recently, however, we have found that activation of the PrL-PVT pathway in STs decreases the incentive value of a food cue and the tendency to sign-track, while inhibition of the same pathway in GTs permits the attribution of incentive value to a food cue and increases the tendency to sign-track [45]. These results suggest that the PrL-PVT pathway acts to suppress the incentive motivational value of a food cue.

In the current study we assessed the role of the PrL-PVT pathway in mediating individual variation in both cue-induced and cocaine-primed reinstatement of drug-seeking behavior. We used an inhibitory DREADD (Designer Receptors Exclusively Activated by Designer Drugs) to selectively “turn off” the PrL-PVT pathway in STs and GTs. We hypothesized that the PrL-PVT pathway acts to suppress the incentive motivational value of the cocaine cue and that chemogenetic inhibition of this pathway would result in disinhibition, thereby, increasing cue-induced reinstatement of drug-seeking behavior selectively in GTs. We also hypothesized that PrL-PVT inhibition would increase cocaine-primed reinstatement of drug-seeking behavior, as the PVT has been shown to play a role in this form of reinstatement as well [41].

In addition to analyzing data within the scope of the ST/GT model, we also analyzed the data within the context of the high-responder (HR)/ low-responder (LR) animal model, which captures individual variation in the acquisition of drug self-administration [46]. In this model, high-responders (HRs) have been repeatedly shown to self-administer psychostimulants at a faster rate compared to low-responders (LRs) [for review see 47]. Despite differences in the acquisition of drug-taking behavior, outbred HRs and LRs do not differ in relapse behavior following psychostimulant use [48]. The objective of this analysis was to determine whether PrL-PVT inhibition similarly affects cue-induced and cocaine-primed drug-seeking behavior in rats using two different models of addiction vulnerability: one that captures individual variation in reinstatement propensity (ST/ GT model); and one that captures individual variation in the acquisition of cocaine self-administration (HR/LR model).

## Methods

All experimental procedures used were approved by the University of Michigan Institutional Animal Care and Use Committee and abided by the standards set in *The Guide for the Care and Use of Laboratory Animals: Eighth Edition*, published by the National Academy of Sciences and revised in 2011. Figure 1a shows the experimental timeline for the procedures described in the following sections. For additional methodological details, see the Supplementary Materials and Methods.

**Figure 1.**
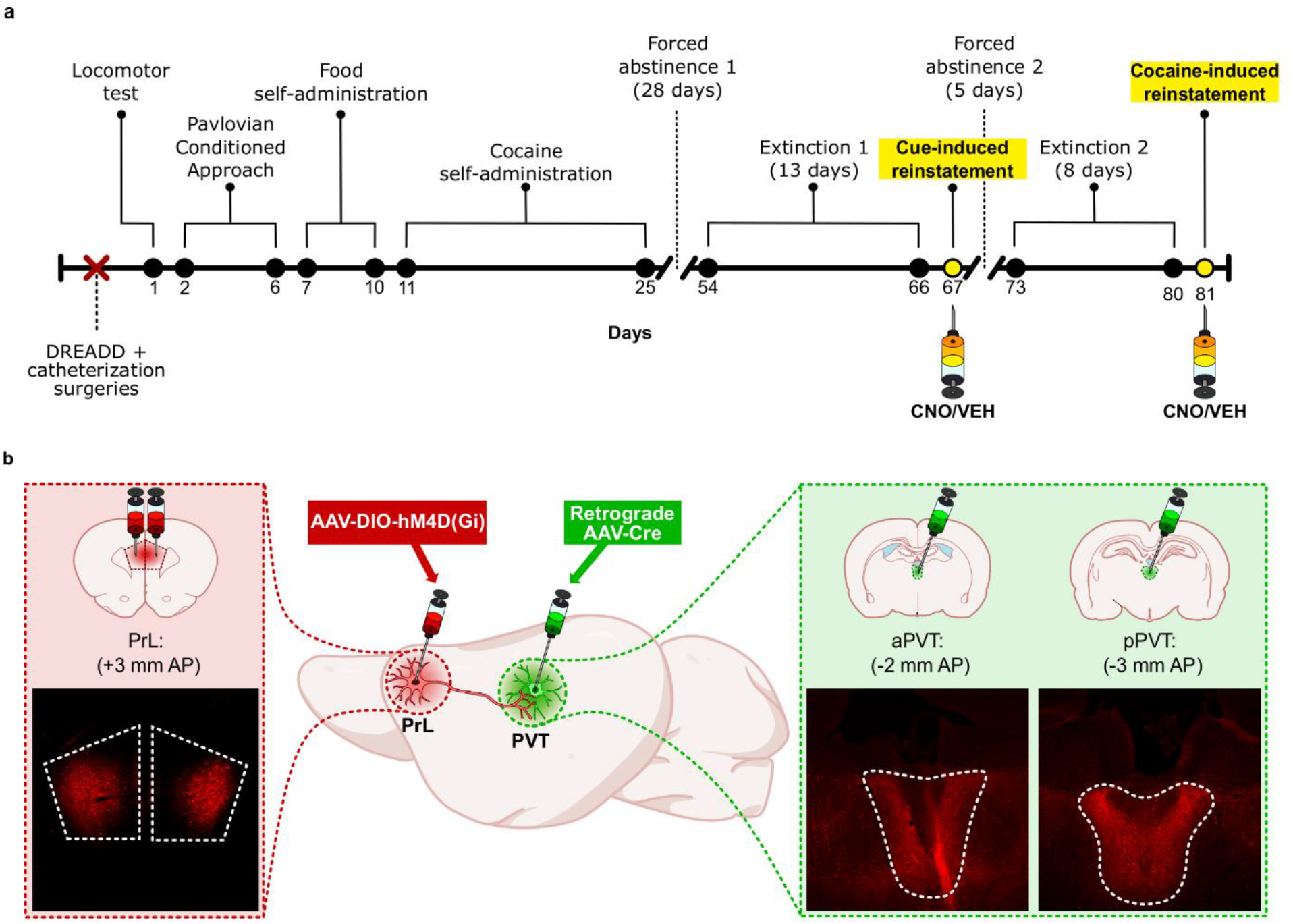
Experimental timeline. **a)** Following recovery from surgery (5 days), rats were characterized based on their locomotor response to a novel environment (i.e. locomotor test) and Pavlovian conditioned approach training. Food self-administration (4 days) and cocaine self-administration (15 days) followed, with a subsequent 28-day forced abstinence period. Extinction sessions occurred for 13 consecutive days prior to the cue-induced reinstatement test. Rats were given an injection (i.p.) of either vehicle (VEH) or 5 mg/kg clozapine-N-oxide (CNO) to activate the G_i_ DREADD prior to the reinstatement test. Following cue-induced reinstatement, rats underwent 5 days of forced abstinence followed by extinction training for 8 consecutive days before a cocaine-primed reinstatement test. Prior to reinstatement, rats were given the same treatment (VEH or CNO) as that prior to the cue-induced reinstatement test, as well as a 15 mg/kg injection of cocaine (i.p.) immediately before being placed into the testing chamber. **b)** Viral infusions into the PrL, aPVT and pPVT for G_i_ DREADD expression in the PrL-PVT pathway. The Cre-dependent Gi DREADD (AAV-DIO-hm4D(Gi)) virus was infused into the PrL, and retrograde AAV-Cre into the anterior and posterior PVT. This resulted in selective DREADD expression, as detected by mCherry in PrL-PVT projection neurons (see representative images; 5x magnification for PrL, 10x magnification for PVT). Abbreviations: AP, anterior posterior (relative to bregma); aPVT, anterior paraventricular nucleus of the thalamus; pPVT, posterior paraventricular nucleus of the thalamus; PrL, prelimbic cortex

### Subjects

A total of 180 (6 cohorts of 30 rats each) male heterogeneous stock (N/NIH-HS) rats bred at the Medical College of Wisconsin were obtained for this study.

### Viruses

The Cre-dependent adeno-associated virus (AAV) carrying an inhibitory DREADD (pAAV-hSyn-DIO-hM4D(Gi)-mCherry, 1.9 × 10^13^ GC/ml, serotype 8) and the retrograde virus carrying Cre (AAV retrograde pmSyn1-EBFP-Cre, 7.6 × 10^12^ GC/ml) were purchased from Addgene (Watertown, MA).

### Drugs

Cocaine HCl (Mallinckrodt, St. Louis, MO) dissolved in 0.9% sterile saline was used in these studies. Clozapine-N-oxide (CNO; Log number 13626-76) was obtained from the National Institute of Mental Health Chemical Synthesis and Drug Supply Program and used at a concentration of 5 mg/ml dissolved in 6% dimethyl sulfoxide (DMSO), which served as the vehicle treatment.

### Surgical procedures

Rats underwent surgery to implant an indwelling jugular catheter for self-administration as previously described [49, 50], followed by stereotaxic surgery for viral infusions (Figure 1b). The G_i_ DREADD was injected bilaterally into the PrL (relative to bregma: AP 3.0, ML: +/− 1.0, DV −4.0) over the course of 5 minutes at a rate of 200 nl/minute, for a total volume of 1 μL. The AAV-Cre was injected into the anterior (relative to bregma: AP −2, ML −1, DV −5.4) and posterior PVT (relative to bregma: AP −3.0, ML −1.0, DV −5.5) over the course of 2 minutes at a rate of 50 nl/minute, for a total of 100 nl per injection site using a 1 μl syringe (see Figure 1a).

### Locomotor test

After 5 days of recovery from surgery, rats underwent a 60-min locomotor test in a novel environment as previously described [51]. Cumulative locomotor movements (vertical and horizontal movements) were calculated and a median split was used to classify rats as either high (HR) or low (LR) responders.

### Pavlovian conditioned approach (PavCA) training

The day following the locomotor test, rats started Pavlovian conditioned approach (PavCA) training as previously described [11]. Briefly, during training, an illuminated lever (conditioned stimulus, CS) entered the chamber for 8-sec, and immediately upon its retraction a food pellet (unconditioned stimulus, US) was delivered into an adjacent food magazine. Lever-CS/ food-US trials were on a variable interval 90-second schedule (range 30-150 seconds). Rats underwent five PavCA training sessions, with one session per day.

Based on the behavior that emerges during Pavlovian training, the PavCA index score [11] was calculated. The PavCA index is a composite score that is generated according to response bias, latency and vigor of responding toward the lever-CS versus the food magazine. Scores range from −1 to 1, with a score of −1 representing individuals with a conditioned response (CR) directed solely towards the food magazine during lever-CS presentation (e.g. goal-tracker, GT), and a score of 1 represents individuals with a CR directed solely towards the lever-CS upon its presentation (e.g. sign-tracker, ST). The PavCA index for session 4 and 5 were averaged to characterize rats into their respective behavioral phenotypes. Rats were characterized as STs if they had a PavCA score between 0.3 and 1, GTs between −0.3 and −1, and intermediate rats (IN), individuals who vacillate between the two CRs, between −0.29 and 0.29.

### Food self-administration

Rats were trained to perform an operant response for food delivery prior to drug self-administration. During these sessions, a nose poke (fixed ratio (FR) 1) into the “active” port resulted in the delivery a single food pellet (same as that delivered during PavCA training), presentation of a cue-light inside the port for 20 seconds, and the house light turning off (i.e. time-out period). In order to minimize location bias, the “active” port was on the opposite side of the food magazine as the lever-CS was during PavCA training. Rats underwent four daily food self-administration sessions.

### Cocaine self-administration

Cocaine self-administration sessions occurred as previously described [44]. Briefly, a nose poke into the active port (FR1) resulted in the intravenous delivery of cocaine, the house light turning off, and presentation of a cue light inside the port for 20-sec. Pokes into the inactive port, and the active port during the 20-sec cue light presentation, were without consequence. Sessions terminated after 3 hours, or once the rat met the infusion criterion (IC) for that session [19–21, 52]. An IC is the maximum number of infusions a rat can receive during a given session. Infusion criterion ensure that rats receive the same number of infusions during each training session, as well as the same number of cocaine cue-light pairings. Rats underwent one cocaine self-administration training session per day for 15 consecutive days. Rats were not moved to the subsequent IC until all had met the IC for at least 2 consecutive days. This resulted in the following schedule: three days at IC5, four days at IC10, 3 days at IC20, and 5 days at IC45. At the conclusion of training, rats underwent a 28-day period of forced abstinence where they were left undisturbed in the colony room.

### Extinction training 1 and 2

During extinction training, nose pokes into both the active and inactive ports were recorded, but without consequence. Sessions lasted for 2 hours, and rats underwent one session per day for a total of 13 sessions (Extinction 1) or 8 sessions (Extinction 2).

### Cue-induced reinstatement test

The test for cue-induced reinstatement occurred the day following the last Extinction 1 training session. Twenty-five minutes prior to the test, rats received either an injection (i.p.) of CNO to activate the G_i_ DREADD, or vehicle. A conditioned reinforcement paradigm was used, such that pokes into the active port resulted in presentation of the drug-associated cue light inside the port and termination of the house light for 5-sec, but no cocaine infusion. The test lasted for 2 h, following which rats underwent five days of forced abstinence prior to Extinction 2 training.

### Cocaine-induced reinstatement test

Following the last Extinction 2 training session, rats underwent a test for cocaine-primed reinstatement. As with the cue-induced reinstatement test, rats received the same treatment (CNO or vehicle) 25-min prior to the session. Before being put into the testing chamber, all rats received a 15 mg/kg (i.p.) injection of cocaine. Pokes into the active and inactive ports were recorded, but without consequence.

### Histology

Once testing was complete, all rats underwent transcardial perfusion and brains were processed for fixation and immunohistochemistry as previously described [32]. Only rats with DREADD expression localized to the PrL and PVT were included in final analysis (See Figure S1). A subset of rats were administered either CNO or vehicle (same as test day treatment) 80 minutes prior to transcardial perfusions so assess treatment-induced Fos expression (see Supplementary Materials and Methods section for methods and results).

### Statistical analyses

For all tests, statistical significance was set at p<0.05. Outliers were identified using the boxplot method [53]. All significant results were subjected to a power analysis and exhibited adequate statistical power (the ability to reject the null hypothesis (1-β) >0.55). Cohen’s d was calculated for each significant pairwise comparison to determine effect size [54]. Data from PavCA training, cocaine self-administration and extinction training sessions were analyzed using a linear mixed-effects model. These analyses were conducted using SPSS Statistics Program (Statistical Package for the Social Sciences), version 22 (IBM, Armok, NY). An ANOVA was used to assess the effects of phenotype and treatment on behavior during the novelty-induced locomotor test and the reinstatement test sessions. A repeated measures ANOVA was used to assess the effects of phenotype and treatment on drug-seeking behavior (i.e. pokes into the active and inactive ports) during the last extinction training session and the reinstatement test sessions. All ANOVAs and correlational analyses were analyzed using StatView, version 5.0 (SAS Institute Inc., Cary, NC, USA). When significant main effects or interactions were detected, post-hoc analyses were conducted using Bonferroni tests to correct for multiple comparisons.

## Results

Rats that successfully completed all training and exhibited accurate DREADD expression (see Figure S1 for detected spread of DREADD expression) comprised the following groups: ST VEH, n=6; ST CNO, n=9; GT VEH, n=16, GT CNO, n=15. The sample size also varies as a function of the number of STs or GTs that we had in a given cohort. IN (n=38) rats were excluded from these studies as this behavioral phenotype was not pertinent to the goals of this study. Refer to the Supplementary Materials and Methods section for additional details regarding excluded rats.

### Behavioral results: ST/GT model

#### Individual differences in PavCA behavior

As expected per their classification, STs and GTs differed in their PavCA index across training (F_4,48_=344.12, p<0.001, Figure 2a) and in their final index scores (F_1,42_=710.34, p<0.001, Figure 2b). There were no significant differences in test day treatment between (Phenotype x Treatment interaction, F_1,42_=0.84, p=0.37) or within phenotypes (effect of Treatment, ST: F_1,13_=0.42, p=0.53; GT: F_1,29_=0.23, p=0.63) on the final PavCA index score (Figure 2b), as experimental groups were counterbalanced according to this metric.

**Figure 2.**
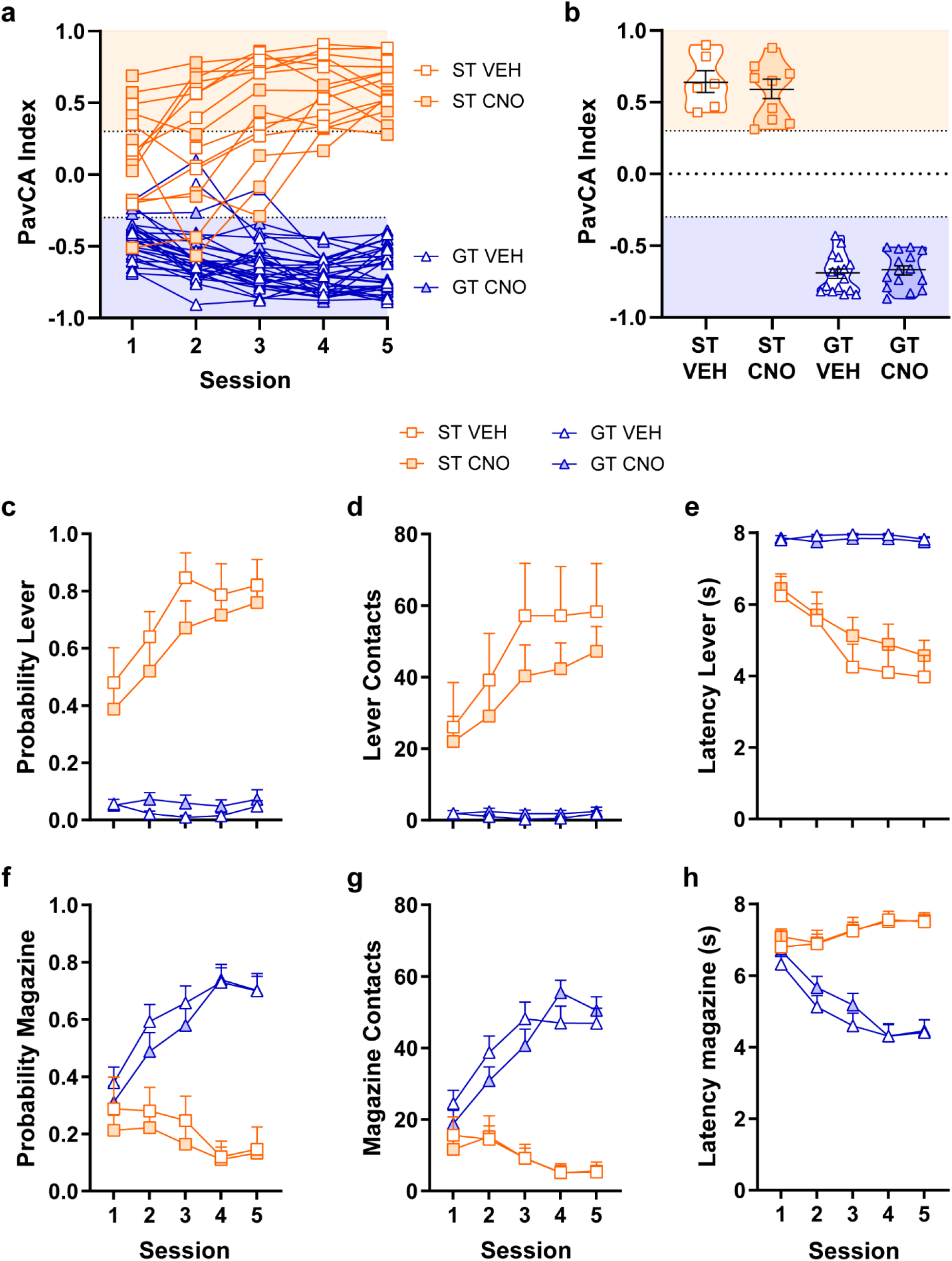
Individual differences in Pavlovian conditioned approach (PavCA) training. **a)** PavCA scores across 5 sessions of training indicated for each individual rat. **b)** Data points illustrate the average PavCA score from session 4 and 5 of training for each individual rat. Mean ± SEM for each group indicated by black bars within violin plots. Scores between 0.3 and 1 indicates a conditioned response directed toward the lever-CS (i.e. ST), whereas a score between −0.3 and −1 corresponds to a conditioned response directed toward the food magazine (i.e. GT). Mean + SEM for **c)** the probability to approach the lever-CS during its presentation, **d)** the number of lever-CS contacts made, **e)** latency to approach the lever-CS, **f)** probability to contact the food magazine during lever-CS presentation, **g)** number of food magazine contacts, and **h)** latency to contact the food magazine during lever-CS presentation. There was a significant effect of Phenotype, Session, and Phenotype x Session interaction for all measures (p<0.01). Sign-trackers (VEH, n=6; CNO, n=9) acquired lever-CS directed behavior, and goal-trackers (VEH, n=16; CNO, n=15) acquired food-magazine directed behavior. Rats did not receive any treatment prior to PavCA training but data are illustrated according to the treatment (i.e. VEH or CNO) received prior to the reinstatement tests.

For each sign-tracking, or lever-directed, measure, there was a significant effect of Session (p<0.0001), Phenotype (p<0.0001) and a Session x Phenotype interaction (p<0.0001) (Figure 2c-e). Compared to GTs, STs showed a greater probability to contact the lever-CS (F_1,42_=85.04, p<0.0001; Figure 2c), greater number of contacts with the lever-CS (F_1,42_=183.67, p<0.0001; Figure 2d), and lower latency to contact the lever-CS (F_1,42_=127.60, p<0.0001; Figure 2e). Post-hoc analyses revealed that STs differed from GTs on all three measures across the five sessions (p<0.0001 for all). There was also a significant effect of Session (p<0.005), Phenotype (p<0.0001) and a Session x Phenotype interaction (p<0.0001) for all measures of goal-tracking, or magazine-directed behaviors (Figure 2f-h). Compared to STs, GTs showed a greater probability to contact the food magazine (F_1,45_=55.81, p<0.0001; Figure 2f), greater number of magazine contacts (F_1,45_=76.83, p<0.001; Figure 2g), and lower latency to approach the food magazine upon lever-CS presentation (F_1,48_=61.62, p<0.0001; Figure 2h). Post-hoc analyses showed that GTs differed from STs on all three measures on sessions two through five (p<0.0001). As expected, there were no significant effects of Treatment between or within phenotypes for lever- or magazine-directed behaviors, as PavCA training occurred prior to treatment. See Table S1 for additional statistical results.

#### STs and GTs that acquired cocaine self-administration did so at the same rate

Acquisition of cocaine self-administration was evident by an increase in nose pokes over the course of training (effect of IC, F_3,85_=84.88, p<0.001), and discrimination between the active and inactive ports (effect of Port, F_1,85_=215.68, p<0.001; Figure 3a). While nose pokes into the active port increased with training (IC x Port interaction, F_3,85_=63.32, p<0.0001), nose pokes into the inactive port did not change (p=0.25; Figure 3a). As expected, there was not a significant effect of Phenotype (F_1,85_=0.60, p=0.44) nor Treatment (F_1,85_=0.80, p=0.78) over the course of training; nor were there significant interactions with these variables. To further assess individual variation in the acquisition of cocaine self-administration, the interinfusion interval (III), or the time (min) between pokes into the active port that resulted in an infusion of cocaine, was analyzed. All rats earned infusions faster as training progressed across IC (effect of IC, F_3,87_= 17.36, p<0.0001), indicating an increase in the motivation to take cocaine (Figure 3b). This measure was not impacted by phenotype (F_1,52_=0.53, p=0.47) or test day treatment (F_1,52_=1.01, p=0.32).

**Figure 3.**
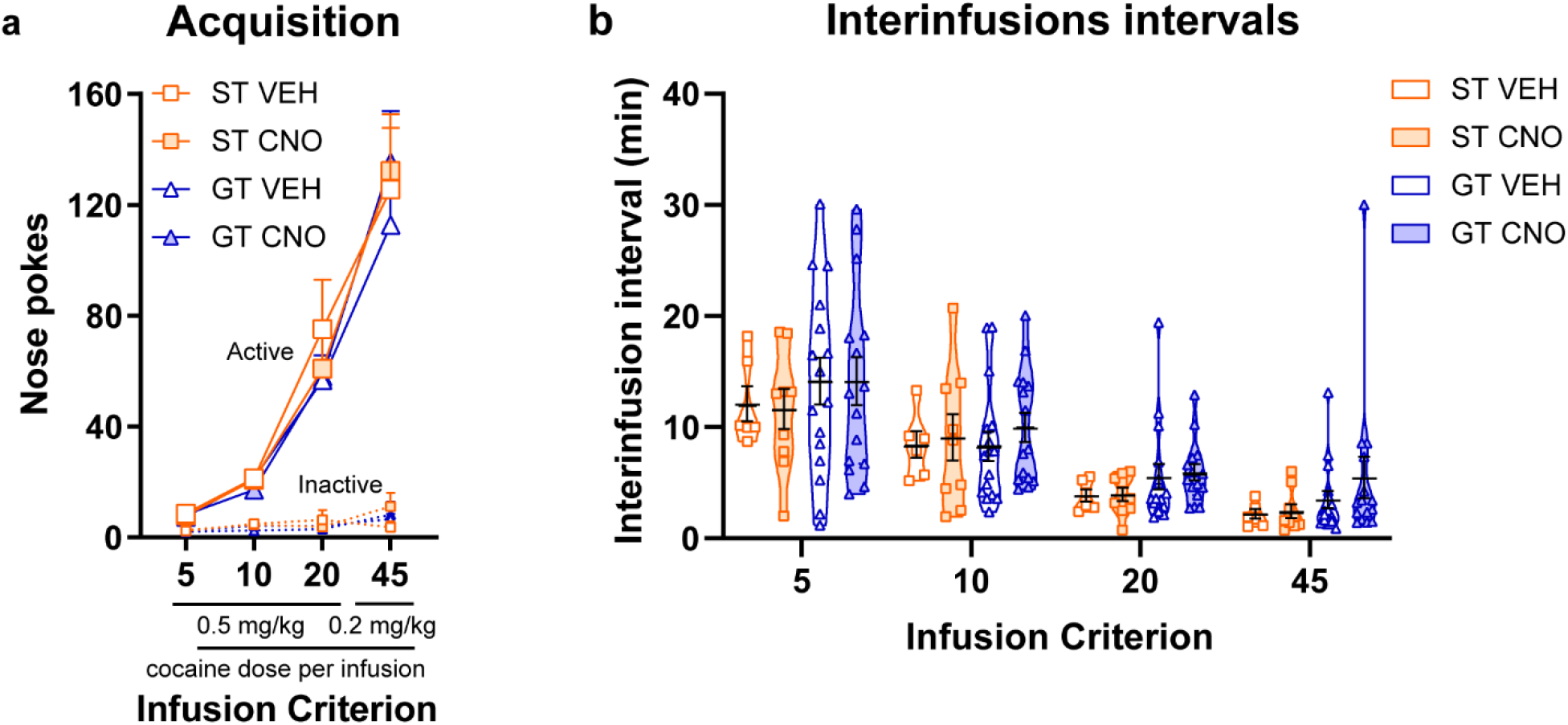
Acquisition of cocaine self-administration in STs and GTs. Mean + SEM for **a)** nose pokes into the active and inactive port for STs and GTs during acquisition of cocaine self-administration across infusion criterion (IC). Rats made more pokes into the active port (p<0.001) across infusion criterion (p<0.001). **b)** Data points illustrate mean interinfusion interval (III) for each rat according to their assigned phenotype (ST, GT) and treatment group (VEH, CNO) at each IC. Mean ± SEM for each group are indicated by black bars within violin plots. The III decreased as training progressed (p<0.001). (ST VEH, n=6, ST CNO, n=9, GT VEH, n=16, GT CNO, n=15)

#### STs and GTs did not differ in extinction of drug-seeking behavior

There were no significant differences between STs and GTs in the extinction of nose-poking behavior (effect of Phenotype, F_1,84_=2.50, p=0.12), and, as expected, test-day treatment did not affect extinction behavior (effect of Treatment, F_1,84_=0.12, p=0.73; Figure 4a). Rats made fewer responses into the ports as training progressed (effect of Session, F_12,84_=24.04, p<0.0001), and continued to differentiate between the two nose ports (effect of Port, F_1,84_=36.77, p<0.0001) until the last few sessions (Session x Port interaction, F_12,84_=5.61, p<0.0001; Session 9, p=0.18; Session 10, p=0.81; Session 11, p=0.07; Figure 4a).

**Figure 4.**
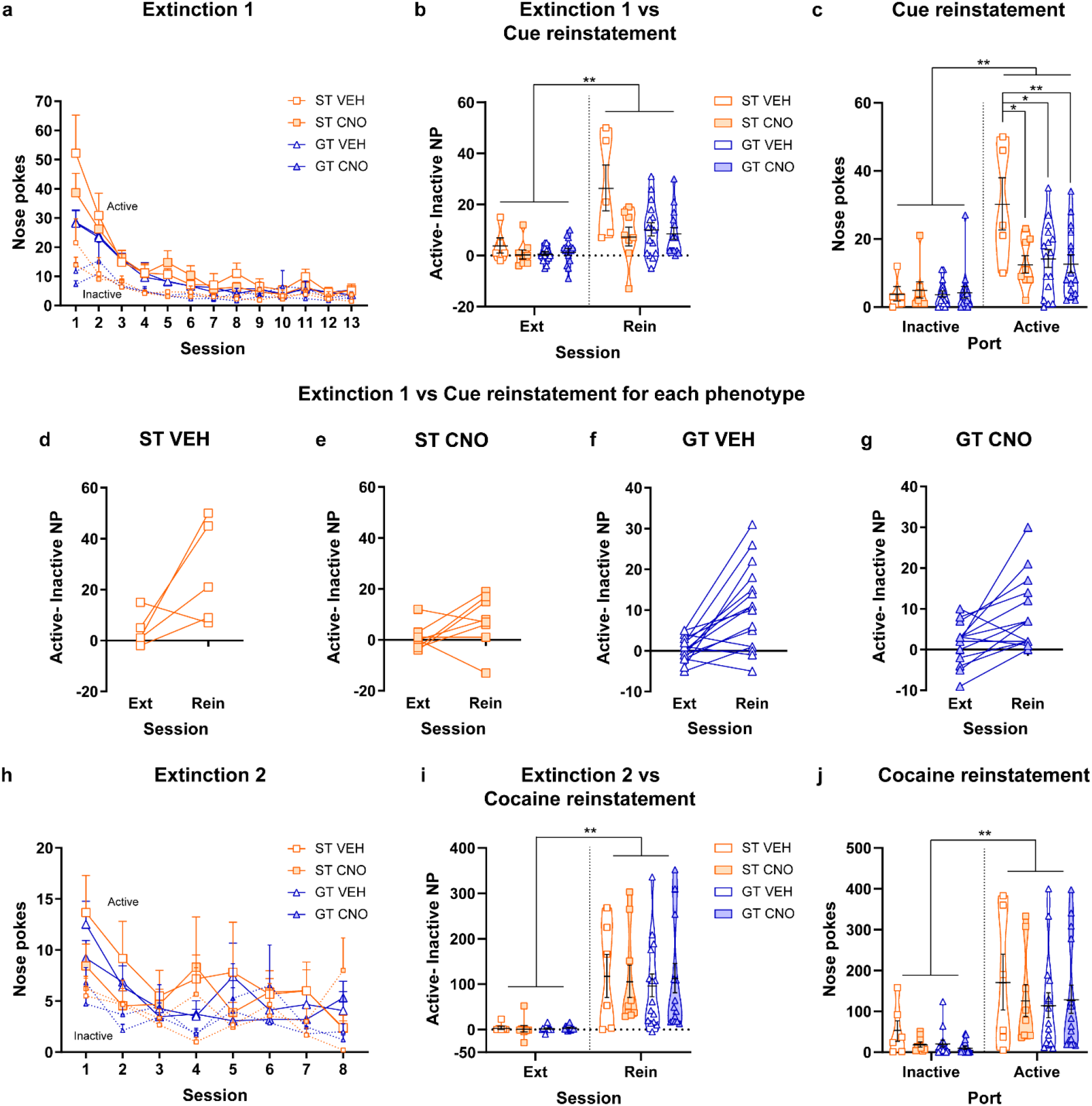
Extinction training and drug-seeking behavior during tests for cue-induced and cocaine-primed reinstatement in STs and GTs. **a)** Mean + SEM for nose pokes (NP) made into the active (solid lines) and inactive (dashed lines) ports across 13 sessions of extinction 1 training in STs and GTs. Rats decreased drug-seeking behavior across training sessions (p<0.0001). **b)** Data points illustrate the mean number of active-inactive NPs made during the last extinction training session compared to the test for cue-induced reinstatement. Black bars within violin plots represent the group mean ± SEM. All rats showed an increase in drug-seeking behavior during the test compared to extinction (p<0.0001). **c)** Individual data points and group mean ± SEM (black bars within violin plots) shown for the number of NPs into the inactive and active port during the test for cue-induced reinstatement. The ST VEH group made more pokes into the active port compared to the GT VEH (p=0.02) and GT CNO (p= 0.01) groups. Inhibition of the PrL-PVT pathway selectively decreased drug-seeking behavior in STs (p=0.02) compared to the ST VEH group. Data shown for each individual rat for the active-inactive NPs made during the last extinction training session (Ext) and the test for reinstatement (Rein) in the **d)** ST VEH, **e)** ST CNO, **f)** GT VEH, and **g)** GT CNO groups. **h)** Mean + SEM number of nose pokes made into the active (solid lines) and inactive (dashed lines) ports across 8 sessions of extinction 2 training. Rats decreased drug-seeking behavior across training sessions (p<0.0001). **i)** Individual data points and mean ± SEM (black bars within violin plots) shown for the number of active-inactive NP made during the last extinction training session compared to the test for cocaine-primed reinstatement. All rats showed an increase in drug-seeking behavior during the test compared to extinction (p<0.0001). **j)** Individual data points and mean ± SEM (black bars within violin plots) shown for the number of nose pokes into the inactive and active port during a test for cocaine-induced reinstatement. All rats showed more responding into the active port relative to the inactive port (p<0.01). *p<0.05, **p<0.01 (ST VEH, n=6 (n=5 for cued reinstatement), ST CNO, n=8, GT VEH, n=16 (n=15 for cocaine reinstatement), GT CNO, n=15)

#### Inhibition of the PrL-PVT pathway selectively decreased cue-induced drug-seeking behavior in STs

All rats exhibited cue-induced reinstatement, as there was a significant increase in drug-seeking behavior (active-inactive NPs) during the test for cue-induced reinstatement compared to the last extinction training session (effect of Session, F_1,40_=33.37, p<0.0001; Figure 4b, d-g; see Supplementary Materials and Methods for additional statistics for active-inactive NPs). During the test for cue-induced reinstatement, more pokes were made into the active port compared to the inactive port (effect of Port, F_1,80_=41.21, p<0.001; Figure 4c). As expected, STs showed enhanced cue-induced drug-seeking behavior relative to GTs (effect of Phenotype, F_1,80_=4.18, p=0.04; Figure 4c) [20, 21]. However, there was a significant effect of Treatment (F_1,80_=4.36, p=0.04) and a Phenotype x Treatment x Port interaction (F_1,80_=4.30, p=0.04). Inhibition of the PrL-PVT pathway rendered STs similar to GTs (Figure 4c). Thus, while STs in the VEH group showed greater responding into the active port compared to both VEH-treated (p=0.02, Cohen’s d=1.14) and CNO-treated GTs (p=0.01, Cohen’s d=1.24), STs treated with CNO did not significantly differ from GTs (GT VEH: p=0.97; GT CNO, p=0.96). CNO-treated STs also exhibited less cue-induced drug-seeking behavior relative to VEH-treated STs (Figure 4c). In STs, inhibition of the PrL-PVT pathway significantly reduced nose-poke responding into the active port (p=0.02, Cohen’s d=1.34), but not the inactive port (p=0.74; Figure 4c). In contrast, in GTs, inhibition of the PrL-PVT pathway did not affect nose pokes into either the active or inactive port in GTs (GTs: CNO vs. VEH, active port, p=0.74; inactive port, p=0.74; Figure 4c). Together, these data suggest that inhibition of the PrL-PVT pathway selectively affects the incentive motivational value of the cocaine-cue in STs. Two rats (ST VEH, n=1; ST CNO, n=1) were excluded from analyses as statistical outliers, however, all statistical outcomes reported remain significant both with or without these outliers.

#### STs and GTs did not differ in extinction of drug-seeking behavior after cue-induced reinstatement

Nose pokes into both ports decreased as training progressed (effect of Session, F_7,135_=6.17, p<0.0001), and did not differ based on phenotype (effect of Phenotype, F_1,87_=1.54, p=0.22) or test day treatment (effect of Treatment, F_1,87_=0.26, p=0.61; Figure 4h; no significant interactions). However, rats continued to differentiate between the ports throughout extinction training (effect of Port, F_1,87_=8.26, p<0.001; Figure 4h). One rat (ST) was excluded from day 8 of analysis as a statistical outlier.

#### Inhibition of the PrL-PVT pathway did not affect cocaine-induced reinstatement of drug-seeking behavior in either STs or GTs

All rats showed greater drug-seeking behavior (active-inactive NPs) during the test for cocaine-induced reinstatement compared to the last extinction training session (effect of Session, F_1,40_=34.48, p<0.0001; Figure 4i; see Supplementary Materials and Methods for additional statistics for active-inactive NPs). During the cocaine-primed reinstatement test, rats made more pokes into the active port compared to the inactive port (effect of Port, F_1,80_=23.13, p<0.0001; Figure 4j). STs and GTs did not differ from one another during this reinstatement test (effect of Phenotype, F_1,80_=0.64, p=0.43) and inhibition of the PrL-PVT pathway did not affect cocaine-primed reinstatement of drug-seeking behavior (effect of Treatment, F_1,80_=1.39, p=0.24; Figure 4j; no significant interactions). Additionally, STs and GTs did not differ in the amount of time spent exhibiting cocaine-induced stereotyped behaviors, suggesting that sensitization to cocaine did not affect reinstatement of drug-seeking behavior (Figure S2). Two rats (GT VEH, n=1; ST CNO, n=1) were excluded from reinstatement analyses as statistical outliers, but there is not a significant effect of Phenotype or Treatment with or without these outliers.

### Behavioral Results: HR/LR model

#### HRs exhibit greater locomotor activity in a novel environment compared to LRs

HRs showed greater locomotor activity in response to a novel environment compared to LRs (effect of Phenotype, F_1,42_=63.02, p<0.0001; Figure 5a), and locomotor activity of rats did not differ between cohorts (effect of Cohort, F_1,38_=1.08, p=0.31; *data not shown*). Additionally, in agreement with prior reports using outbred populations [12], there was not a significant correlation between novelty-induced locomotor activity and PavCA index for STs (r^2^=0.10, p=0.25) or GTs (r^2^=0.01, p=0.69), indicating that these two traits – “sensation-seeking” and the propensity to attribute incentive salience to reward cues – are independent from one another.

**Figure 5.**
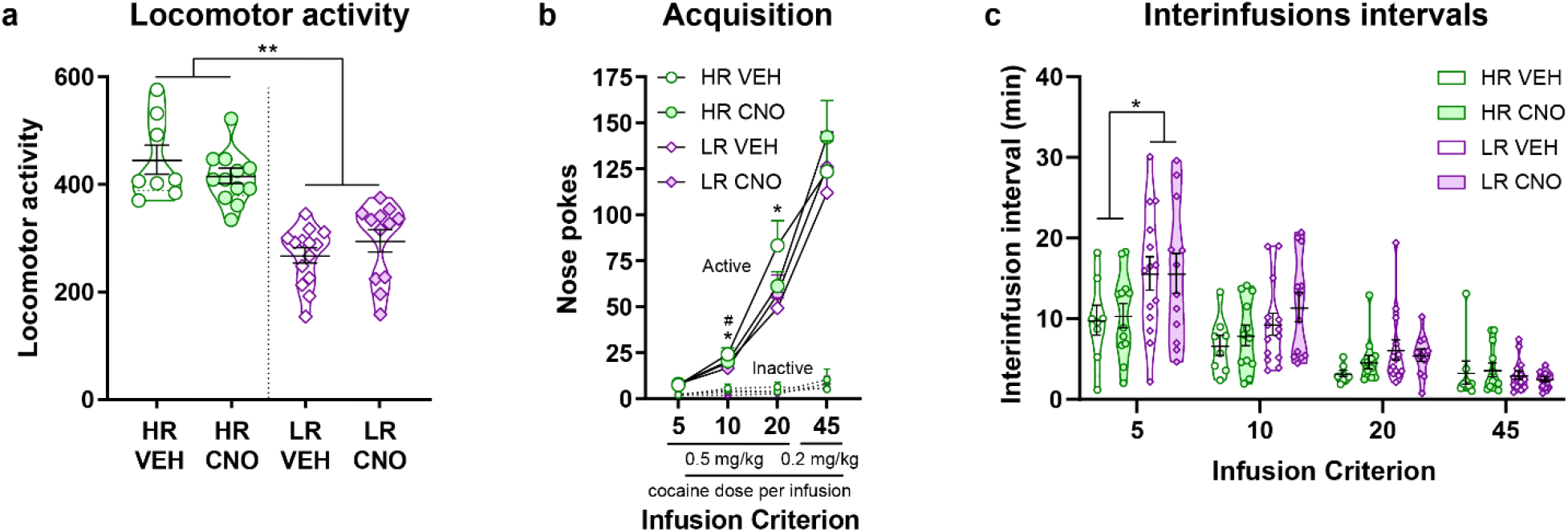
Locomotor response in a novel environment and acquisition of cocaine self-administration. **a)** Individual data points represent the mean cumulative locomotor activity in a novel environment. Black bars within the violin plots represent the mean ±SEM for each group. HRs show greater locomotor activity compared to LRs (effect of Phenotype, p<0.0001). Rats did not receive treatment prior to the locomotor test. Mean + SEM for **b)** nose pokes made into the inactive and active port across infusion criterion (IC). Rats made more pokes into the active port (p<0.0001) across infusion criterion (p<0.0001). HRs showed more drug-seeking behavior during IC10 (p=0.01) and IC20 than LRs (p=0.02). Rats in the vehicle-treated group also showed more drug-seeking behavior during IC10 (p=0.04) compared to rats treated with CNO. **c)** Individual data points representing the mean interinfusion interval (III) for HRs and LRs at each IC. Black bars within violin plots represent the mean ± SEM. III decreased across all groups as training progressed (p<0.001). However, HRs earned infusions faster than LRs at IC5 (p=0.01), and there was a trend toward HRs continuing to earn infusions faster than LRs at IC 10 (p=0.06) and IC20 (p=0.07). *p<0.05 for HR compared to LR, ^#^p<0.05 for vehicle-treated versus CNO-treated rats (HR VEH, n=8, HR CNO, n=12, LR VEH, n=14, LR CNO, n=12)

#### HRs and LRs differed in the acquisition of cocaine self-administration

All rats acquired cocaine self-administration (effect of IC, F_3,85_=103.23, p<0.0001) and discriminated between the two ports (effect of Port, F_1,85_=251.14, p<0.0001; Figure 5b). Responses in the ports were dependent upon the IC (IC x Port interaction, F_3,85_=76.64, p<0.0001), such that rats increased nose pokes made into the active port as training progressed (p<0.0001), but not the inactive port. Behavior in HRs and LRs differed with each successive IC (Phenotype x IC interaction, F_3,85_=4.07, p=0.01) and, compared to LRs, HRs showed greater nose-poking behavior during IC10 (p=0.01, Cohen’s d=0.30) and IC20 (p=0.02, Cohen’s d=0.26; Figure 5b). A significant Treatment x IC interaction (F_3,85_=3.35, p=0.02) was also present, revealing that rats in the vehicle-treatment group showed greater nose-poking behavior at IC10 (p=0.04, Cohen’s d=0.19) than rats in the CNO treatment group. Notably, this effect size is small, and the two treatment groups did not differ at IC5 (p=0.67), IC20 (p=0.32) or IC45 (p=0.26). Further, there was not a significant Treatment x Port x IC interaction (F_3,85_=1.30, p=0.25).

While all rats decreased their III as training progressed (effect of IC, F_3,74_=30.30, p<0.0001), HRs self-administered cocaine at a faster rate than LRs across infusion criterion (effect of Phenotype, F_1,50_=6.78, p=0.01; Figure 5c). This difference was most apparent during IC5, (Phenotype x IC interaction, F_3,74_=4.16, p=0.01; at IC5, p=0.01, Cohen’s d=0.82), and there were no significant differences during IC10 (p=0.06), IC20 (p=0.07) and IC45 (p=0.31) (Figure 5c). Thus, despite the fact that this paradigm controlled for the maximum number of infusions received, we were able to capture individual differences in the acquisition of drug-taking behavior in outbred HR/LR rats.

#### HRs and LRs did not differ in extinction of drug-seeking behavior

All rats decreased nose pokes into both ports (effect of Session, F_12,84_=26.67, p<0.001) and differentiated between the two ports across extinction training sessions (effect of Port, F_1,84_=41.94, p<0.0001; Figure 6a). As training progressed rats discriminated less between the two ports (Session x Port interaction, F_12,84_=6.19, p<0.0001), with no significant differences between the ports during session 9 (p=0.12), 10 (p=0.93), and 11 (p=0.07). There was not a significant main effect of Phenotype (F_1,84_=3.00, p=0.09), but nose-poking behavior differed between HRs and LRs depending upon impending test day treatment (Phenotype x Treatment interaction, F_1,84_=11.54, p<0.001). Rats in the vehicle groups made more pokes into both ports compared to rats in the CNO groups (p=0.001; Figure 6a). Test day treatment also affected responding into the active and inactive ports within each phenotype (HR, p=0.01, Cohen’s d=0.3; LR, p=0.04, Cohen’s d=0.23; Figure 6a). Importantly, there were no significant differences between HRs and LRs (effect of Phenotype, F_1,84_=2.20, p=0.14) or test day treatment (effect of treatment, F_1,84_=1.97, p=0.16) on nose pokes made into either port during the last extinction training session. Thus, by the conclusion of extinction training, all rats were behaving the same.

**Figure 6.**
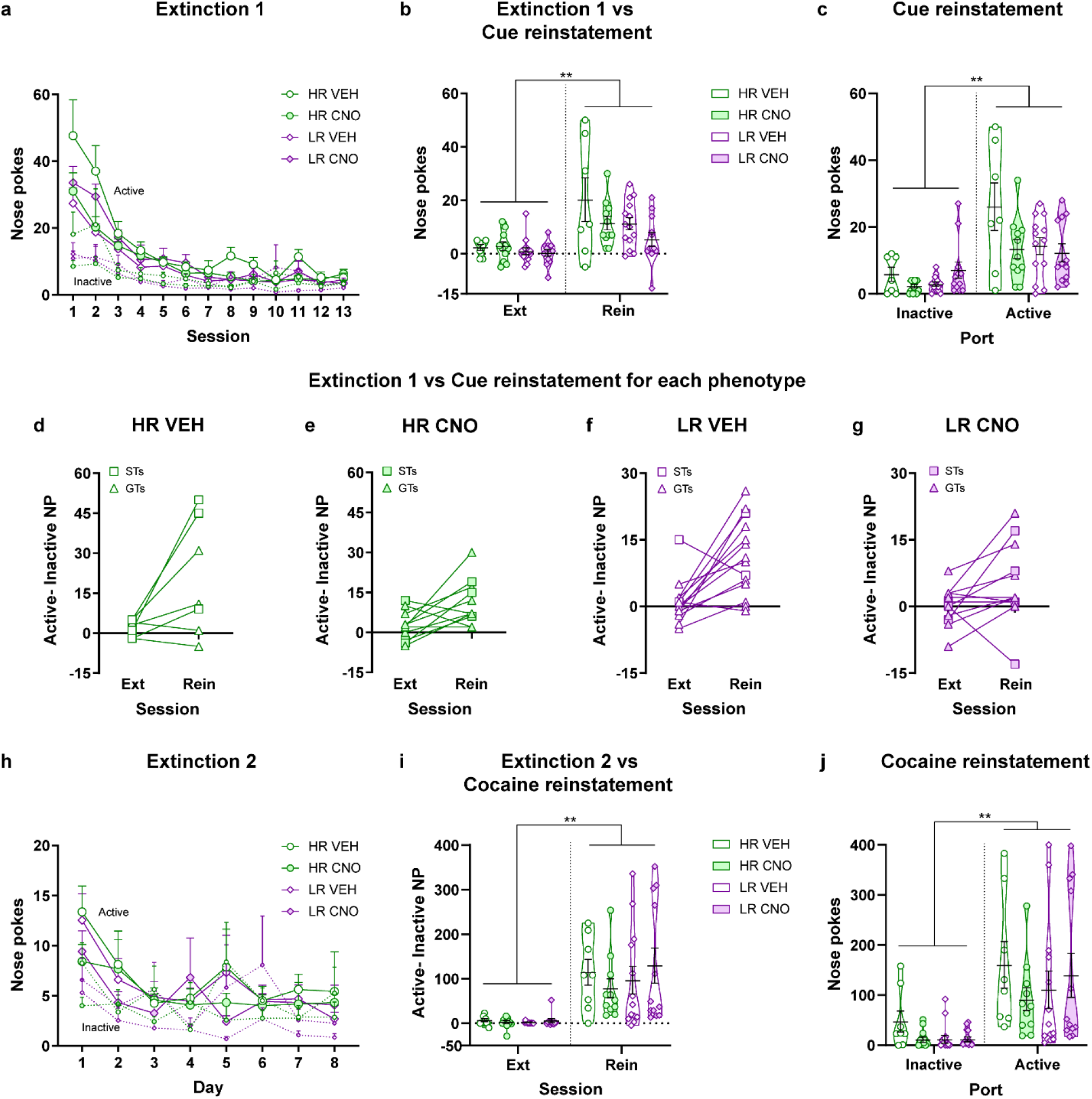
Extinctions and drug-seeking behavior during tests for cue-induced and cocaine-primed reinstatement in HRs and LRs. **a)** Mean + SEM for nose pokes (NP) made into the active (solid lines) and inactive (dashed lines) ports across 13 sessions of extinction training in HRs and LRs. Rats decreased drug-seeking behavior across training sessions (p<0.0001). Rats in the HR vehicle group showed greater drug-seeking behavior throughout training compared to rats in the LR vehicle group (p=0.001) and HR CNO group (p=0.01). LR rats in the CNO group also showed greater drug-seeking behavior compared to rats in the LR vehicle group (p=0.04). **b)** Individual data points for the number of active-inactive NPs made during the last extinction training session compared to the test for cue-induced reinstatement. Black bars within violin plots represent the mean ± SEM for each group. All rats showed an increase in drug-seeking behavior during the test compared to extinction (p<0.0001). **c)** Individual data points for the number of nose pokes made into the inactive and active port during the test for cue-induced reinstatement. Black bars within violin plots represent the mean ± SEM for each group. Data shown for each individual rat for the active-inactive NPs made during the last extinction training session (Ext) and the test for reinstatement (Rein) in the **d)** HR VEH, **e)** HR CNO, **f)** LR VEH, and **g)** LR CNO groups. The symbol of the individual data point indicates if the rat was classified as a ST or GT based on their behavior during PavCA training. **h)** Mean + SEM number of nose pokes made into the inactive and active port across 8 sessions of extinction training. All rats decreased drug-seeking behavior across training sessions (p<0.0001). **i)** Individual data points for the number of active-inactive NPs made during the last extinction training session compared to the test for cocaine-primed reinstatement. Black bars within violin plots represent the mean ± SEM for each group. All rats responded more during the reinstatement test relative to the last extinction session (p<0.0001). **j)** Individual data points for the number of nose pokes made into the inactive and active port during the test for cocaine-primed reinstatement. Black bars within violin plots represent the mean ± SEM for each group. All rats responded more into the active port relative to the inactive port (p<0.0001) (HR VEH, n=8 (n=7 for cue-induced reinstatement); HR CNO, n=11; LR VEH, n= 14 (n=13 for cocaine-primed reinstatement), LR CNO, n= 12) **p<0.01

#### Inhibition of the PrL-PVT pathway did not affect cue-induced drug-seeking behavior in HRs or LRs

All rats exhibited cue-induced reinstatement, as there was a significant increase in drug-seeking behavior (active-inactive NPs) during the test for cue-induced reinstatement compared to the last extinction training session (effect of Session, F_1,40_=31.89, p<0.0001; Figure 6b, d-g; see Supplementary Materials and Methods for additional statistics for active-inactive NPs). During the test for cue-induced reinstatement, rats made more pokes into the active port compared to the inactive port (effect of Port, F_1,80_=38.84, p<0.0001; Figure 6c). CNO administration differentially affected nose-poke responding for HRs and LRs relative to their vehicle-treated controls (Phenotype x Treatment interaction, F_1,80_=6.05, p=0.02; Figure 6c); and HRs and LRs differed in their responding across ports (Phenotype x Port interaction, F_1,80_=3.99, p<0.05) during the test session. However, there were no significant post-hoc analyses nor a significant Phenotype x Treatment x Port interaction (F_1,80_=0.16, p=0.69). Two rats (HR VEH, n=1, HR CNO, n=1) were excluded from the analyses as statistical outliers, but there was not a significant Phenotype x Treatment x Port interaction even with the outliers included (F_1,84_=1.15, p=0.29).

#### HRs and LRs did not differ in extinction of drug-seeking behavior after cue-induced reinstatement

Pokes into both ports decreased (effect of Session, F_7,84_=9.82, p<0.0001) and rats discriminated between both ports across extinction training (effect of Port, F_1,84_=6.13, p=0.02; Figure 6h) following the cue-induced reinstatement test. HRs and LRs did not differ from one another (effect of Phenotype, F_1,84_=0.76, p=0.39), and test day treatment did not affect behavior during extinction training (effect of Treatment, F_1,84_=0.12, p=0.73; Figure 6h).

#### Inhibition of the PrL-PVT pathway did not affect cocaine-induced reinstatement of drug-seeking behavior in either HRs or LRs

All rats exhibited greater drug-seeking behavior during the cocaine-primed reinstatement test compared to the last extinction training session (effect of Session, F_1,40_=39.78, p<0.0001; Figure 6i). During the test, both HRs and LRs preferred the active port relative to the inactive port (effect of Port, F_1,80_=26.24, p<0.0001; Figure 6j). There were no significant difference between phenotypes (effect of Phenotype, F_1,80_=0.13, p=0.72) and no effect of Treatment (F_1,80_=0.83, p=0.37). There were also no significant interactions, indicating that inhibition of the PrL-PVT pathway did not affect cocaine-induced reinstatement of drug-seeking behavior in HRs or LRs. Lastly, HRs and LRs did not differ in the amount of time spent exhibiting cocaine-induced stereotyped behaviors (Figure S4).

## Discussion

While the PrL-PVT pathway has been implicated in cue-reward learning [45, 55–57], the present study assessed the role of this pathway in mediating individual variation in cue-induced and cocaine-primed reinstatement of drug-seeking behavior. Sign-tracker and goal-tracker rats, categorized based on their propensity to attribute incentive salience to reward-paired cues, did not differ in behavior during cocaine self-administration [19–21, 44] or extinction training prior to the reinstatement tests [19, 22, 44, 58, 59]. Consistent with prior studies, however, STs showed greater cue-induced drug-seeking behavior relative to GTs [20, 21]. Notably, chemogenetic inhibition of the PrL-PVT pathway selectively decreased cue-induced drug-seeking behavior in STs, eliminating phenotypic differences. When the same rats were categorized as high- or low-responders, according to their locomotor response to a novel environment, PrL-PVT inhibition had no effect on drug-seeking behavior. Thus, the PrL-PVT pathway appears to play a selective role in cue-induced reinstatement in individuals with a greater propensity to attribute incentive salience to reward cues.

Unlike prior studies [19], we did not observe differences between STs and GTs in cocaine-primed reinstatement of drug-seeking behavior. While we used the same drug dose as that used in previous experiments [19], methodological differences may have contributed to these seemingly discrepant findings. In the current study, rats did not receive cocaine for approximately 2 months prior to the cocaine-primed reinstatement test; while in the study conducted by Saunders and Robinson (2011) rats were abstinent for only 1 week [19]. It is possible that the presence of the prolonged forced abstinence period in the current study, which is known to result in cellular adaptations following cocaine experience [for review see 60], eliminated phenotypic differences. In relation, it is possible that the dose of cocaine used in the present study masked individual differences in drug-seeking behavior between STs and GTs that a lower dose might have revealed [61]. Nonetheless, under these experimental conditions, the PrL-PVT pathway does not appear to mediate cocaine-induced drug-seeking behavior.

The reported results challenge our hypothesis that the PrL-PVT pathway inhibits the incentive motivational value of reward cues. This hypothesis is based on our previous work that assessed the role of this pathway during Pavlovian learning [45], whereas the current study utilized an instrumental procedure. The neural circuitry mediating these two forms of associative learning are different [62–65]; thus, it is possible that the PrL-PVT pathway plays a distinct role in each of these forms of learning. Additionally, Campus et al. used a natural reward (e.g. food) [45], whereas here we used cocaine, which can induce more substantial neuroplastic changes within the motive circuit [for review see 66, 67–69], especially during a period of prolonged abstinence following drug exposure [for review see 70]. It is possible, therefore, that cocaine-induced neuroplastic changes uniquely alter the PrL-PVT pathway and its functional role. In support, cells in the PrL and posterior PVT become active shortly after the conclusion of cocaine self-administration training [57], and inhibiting this pathway at this time reduces cue-induced drug-seeking behavior [57]. Based on the current findings, we postulate that the effects of cocaine on neuronal plasticity within the PrL-PVT pathway are dependent on one’s inherent tendency to attribute incentive motivational value to reward cues.

We speculate that inhibition of the PrL-PVT pathway differentially affects cue-induced drug-seeking behavior in STs and GTs by way of regulating the downstream projection to the NAc. The PVT-NAc pathway has been shown to encode food cue-reward relationships [71], and we recently showed that manipulation of the PrL-PVT pathway affects dopamine release in the NAc and does so differently in STs vs. GTs [45]. PVT terminals to the NAc terminate in close proximity to dopamine neurons and can elicit dopamine release [34], and NAc dopamine contributes to greater cue-induced drug-seeking behavior in STs relative to GTs [21]. Further, the PVT-NAc pathway has been shown to mediate cocaine, but not sucrose, self-administration and undergoes cocaine-induced neuroplastic changes following prolonged forced abstinence resulting in an increase in synaptic strength [72]. Thus, it is possible that inhibition of PrL-PVT projection neurons affects downstream communication to the NAc and attenuates dopamine release and, correspondingly, cue-induced drug-seeking behavior in STs. The role of the PrL-PVT pathway in incentive motivational processes and, thereby, cue-induced drug-seeking behavior, is likely mediated via this final common path between the PVT and the NAc.

While STs are more vulnerable to cue-induced relapse, GTs show greater drug-seeking behavior in response to both discriminative stimuli [24] and contexts [22] associated with drug taking. It is perhaps not surprising, therefore, that a pathway that appears to mediate the incentive motivational value of a cocaine-cue does not affect cue-induced drug-seeking behavior in GTs. Indeed, it is possible that, in GTs, the PrL-PVT pathway is selectively involved in context-induced reinstatement. The incentive motivational value of contextual cues in GTs relies, at least in part, on signaling within the prefrontal cortex [24], and the dorsomedial prefrontal cortex (including the PrL) is necessary for context-induced cocaine-seeking behavior [73]. Furthermore, the PVT is recruited following re-exposure to a context previously paired with cocaine [74, 75] as well as a test for context-induced reinstatement [cocaine 76, ethanol 77]; and decreasing neuronal activity in the PVT attenuates cocaine conditioned place preference [78] and context-induced reinstatement of ethanol-seeking behavior [40]. These findings support a potential role for the PrL-PVT pathway in context-mediated drug-seeking behavior. Ongoing studies are being conducted to determine whether this pathway is differentially involved in context-induced reinstatement in GTs and STs.

In summary, these results highlight the complex role of cortical projections from the PrL to the PVT in cue-reward learning. We report here that PrL-PVT inhibition selectively attenuates cue-induced drug-seeking behavior in rats with high relapse propensity (STs), but does not affect cocaine-primed reinstatement of drug-seeking behavior. Thus, this pathway plays a role in encoding the incentive value of a cocaine-cue, but not the motivational value of the interoceptive effects of the drug. Drug-induced neuroplastic changes both within the PrL-PVT pathway and the downstream PVT projection to the NAc likely contributes to these effects, and will be the focus of future investigations.

## Acknowledgements

The experiments were designed by BNK and SBF, and conducted by BNK, PC, MSK, SEC and AGI. Data was analyzed by BNK, and BNK and SBF wrote the manuscript. We would like to thank Dr. Leah C. Solberg Woods and Katie Holl (both Medical College of Wisconsin) for maintaining the heterogeneous stock rat colony used in these studies, and Maurice Chojecki and Erin Canfield (both University of Michigan) for their technical assistance during this study. We would also like to thank Dr. Joshua Haight for his helpful insight into the histological analyses performed for these studies, and Drs. Aram Parsegian and Ignacio R. Covelo for their feedback on a previous version of this manuscript.

## Supplementary materials and methods

### Methods

The following are detailed methods to supplement those included in the main text.

#### Subjects

Throughout the study, rats had ad libitum access to food and water and were housed in a climate-controlled room with a 12-hour light: dark cycle (lights came on at either 06:00 h or 07:00 h depending upon daylight savings time). Upon arrival, rats were pair-housed until surgeries, after which they were single-housed for the remainder of the study to protect the catheters from possible damage from a cage mate. All behavioral training occurred during the light cycle, between 08:00 h and 18:00 h, with specific testing times for each procedure included below.

#### Surgical procedures

After arrival in our housing facilities, rats were allowed to acclimate to the housing room for a minimum of 5 days prior to surgery. Rats were anesthetized using ketamine (90 mg/kg i.p.) and xylazine (10 mg/kg i.p.) and implanted with the jugular vein catheter. After the surgery was completed, rats were given a 5 ml injection (s.c.) of saline to minimize dehydration and facilitate recovery. To ensure sufficient recovery, rats underwent stereotaxic surgery 24 h to 48 h after catheterization surgery.

For stereotaxic surgery, rats were anesthetized using 5% isoflurane, and then maintained under 2% isoflurane for the duration of the surgery. Each stereotaxic apparatus (David Kopf Instruments, Tujunga, CA) was outfitted with two digital manipulator arms (Stoelting, Wood Dale, IL). Rats were fitted into the ear bars, and then the scalp was cleaned with a series of Betadine solution (Purdue Products, Stamford, CT) and ethanol wipes prior to an incision being made to expose the skull. Bregma was determined, and the skull was then leveled such that lambda was within +/− 0.1 mm of the bregma coordinates. The G_i_ DREADD was injected bilaterally into the PrL using a standard infusion pump (Pump 11 Elite, Harvard Apparatus) to depress a 5 μl Hamilton syringe with P50 tubing connecting it to a 31-gauge injector. The AAV-Cre was injected into the anterior and posterior PVT using a 1 μl Hamilton Microsyringe. Injections into the PVT were made at a 10° angle to the midline to avoid puncturing the superior sagittal sinus and causing unnecessary bleeding. At the completion of each injection, the injector was kept in place for an additional five minutes to allow the virus to diffuse away from the injector and into the tissue. At the conclusion of the surgery, surgical staples were used to bring the scalp back together.

Prior to the start of each surgery, and 24 h following surgery, an injection of 5 mg/ml of Carprofen was given as an analgesic. Additionally, rats received daily infusions i.v. of heparin (100 units/ml, 0.05 ml) and gentamicin sulfate (1 mg/ml, 0.05 ml) in order to decrease the chance of infection and maintain catheter patency until the conclusion of cocaine self-administration training. Following surgeries, rats were given a minimum of five days of recovery prior to the start of behavioral training. All surgical staples and sutures were removed within ten days of surgeries. Prior to the start of cocaine self-administration, catheter patency was checked using methohexital sodium diluted in sterile saline (10 mg/ml i.v., 0.1 ml). Rats were removed from the study due to lack of patency if the rat did not become ataxic within 10 seconds of infusion. Additionally, if rats stopped self-administering cocaine in the midst of the study, catheter patency was checked, and rats removed from the study accordingly.

#### Locomotor test

After the surgical recovery period, rats underwent a 60-min locomotor test. Testing chambers (43 × 21.5 cm floor area, 25.5 cm high) were outfitted with infrared beams to track both lateral (beams 2.3 cm above grid floor) and rearing (beams 6.5 cm from grid floor) movements. All testing occurred between 10:00 h and 16:00 h under red light. Lateral and rearing movements were recorded in 5-min increments. At the conclusion of the test, rats were returned to their home cages in the colony room. Cumulative locomotor movements (i.e. rearing and lateral movements) were used to classify rats into their respective phenotypes. Rats for this study were run in six separate cohorts and classified as HRs and LRs within each cohort using a median split based on the cumulative locomotor movements.

#### Pavlovian conditioned approach (PavCA) training

Two days prior to the start of PavCA training rats were given 45-mg banana-flavored grain pellets (about 25 pellets per rat; Bio-Serv, Flemington, NJ, USA) in their home cage to habituate them to the food reward used during training. Rats started PavCA training the day following the locomotor test. Standard behavioral testing chambers (MED Associates, St. Albans, VT, USA; 20.5 × 24.1 cm floor area, 29.2 cm high) were used. Chambers were situated in a sound-attenuating boxes fitted with a fan to provide constant background noise and air circulation. Within each chamber, a food magazine was situated 6-cm above the flooring and attached to a pellet dispenser. When the photo beam within the food magazine was broken a “contact” to the magazine was recorded. A retractable lever that illuminated upon presentation was situated to either the right or left of the food magazine and at the same height. In order for a lever “contact” to be registered and recorded, a minimum of 10-g of force had to be exerted. A white house light in the center of the opposite wall (1-cm from the top of the chamber) remained illuminated for the duration of each session.

Rats underwent two days of pre-training, with one session a day. During pre-training, the food magazine was baited with two banana-flavored grain pellets to direct the rats’ attention to the location of reward delivery. After a 5-min acclimation period, the house light turned on and the session started, lasting approximately 12.5 minutes. During pre-training sessions, the lever remained retracted, and pellets were delivered to the food magazine on a variable interval 30-second schedule (range 0-30 seconds). Sessions consisted of 25 trials, with one pellet being delivered per trial. At the conclusion of the session, the house light turned off. The number of food magazine entries made and pellets remaining in the food magazine were recorded at the end of each session.

Following pre-training, rats underwent five PavCA training sessions, with one session per day. As with pre-training, the start of each session was signaled with the house light turning on, and the conclusion with the house light turning off. Each training session consisted of 25 lever-CS/ food-US trials on a variable interval 90-second schedule (range 30-150 seconds). Each session lasted approximately 40-min, and all training occurred between 10:00 h and 16:00 h.

For each PavCA training session, Med Associates software recorded the following information: (1) number of magazine contacts made during the 8-sec lever-CS presentation, (2) latency to the first magazine contact during lever-CS presentation, (3) number of lever-CS contacts, (4) latency to the first lever-CS contact, (5) the number of magazine entries made during the inter-trial interval (i.e. contacts made between lever-CS presentations). Using these measures, the PavCA index was calculated for each rat and used to characterize rats into their behavioral phenotypes based on the CR that emerged during training. The following formula was used to calculate the PavCA index: [Probability Difference Score + Response Bias Score + (−Latency Difference Score)/3] [11].

#### Food self-administration

Following PavCA training, rats had a 24-hour break during which they remained in the colony room undisturbed before starting food self-administration training. Food self-administration took place in the same testing chambers as PavCA training; however, the chambers were reconfigured such that the lever was removed, and two nose ports were put on either side of the food magazine. The house light and a discrete cue light within the active port turned on one minute after the program was started. The cue light remained on to direct the rat’s attention to the port and was turned off following the first poke into the active port. Pokes into the active port resulted in the delivery of one 45-mg banana flavored grain pellet (FR1 schedule of reinforcement), the same reward used during PavCA training, as well as presentation of the cue-light inside the port for 20 seconds. Additional pokes made into the active port during the 20-sec cue-light presentation period were recorded but without consequence. During cue-light presentation, the house light also turned off. Pokes into the inactive port were recorded, but without consequence. Rats underwent four days of food self-administration testing, with one session per day. Sessions were terminated after rats had acquired 25 pellets, or after 30-min. All training occurred between 10:00 h and 14:00 h.

#### Cocaine self-administration

At the conclusion of food self-administration, rats were left undisturbed in the colony room for 24 hours prior to the start of cocaine self-administration. Testing chambers remained in the same configuration as food self-administration, with the exception that the food magazine and pellet dispenser were removed. The house light and cue light in the active port turned on one minute after the session started. In contrast to food self-administration, the cue light remained on for 20-sec at the start of the session, during which time a poke into the active port could result in an infusion of cocaine. Throughout the session, pokes into the active port (FR1 schedule of reinforcement) resulted in the presentation of the cue light for 20-sec and a 0.5 mg/kg infusion of cocaine (Mallinckrodt, St. Louis, MO) diluted in 0.9% sterile saline, delivered in 25 μl over 1.6 seconds. However, during the IC45 sessions, the cocaine infusion dose was decreased to 0.2 mg/kg/infusion to enhance drug-seeking behavior. Additional pokes into the active port during the 20-sec cue light presentation were recorded, but without consequence. Pokes made into the inactive port were recorded, but also without consequence. Rats underwent one cocaine self-administration training session per day for 15 consecutive days between 8:00 h and 18:00 h. In order to move to the next IC and remain in the study, rats had to meet the IC for two consecutive sessions and maintain catheter patency. At the conclusion of training, rats underwent a 28-day period of forced abstinence where they were left undisturbed in the colony room.

#### Extinction training 1 and 2

Testing chambers remained in the same configuration as that during cocaine self-administration, however pokes into both the active and inactive nose port were recorded, but without consequence (i.e. pokes into the active port did not result in cue-light presentation or drug delivery). Sessions lasted for 2 hours, and rats underwent one session per day for a total of 13 sessions (Extinction 1) or 8 sessions (Extinction 2). Rats had to make less than ten pokes into the active port for two consecutive sessions in order to be eligible to move to the subsequent reinstatement test, or they were eliminated from the study. All training took place between 09:00 h and 18:00 h. After the last training session, rats were moved to an adjacent testing room and given an i.p. injection of 6% DMSO in 0.9% sterile saline (CNO vehicle) to habituate them to the injections that would occur prior to the reinstatement test.

#### Cue-induced reinstatement test

Rats underwent a test for cue-induced reinstatement the day following the last extinction training 1 session (i.e. session 13). Rats were counterbalanced into treatment groups based on behavior during PavCA training (lever- and magazine-directed behavior, PavCA index), locomotor index (cumulative locomotor movements), and behavior during cocaine self-administration and extinction training (nose pokes made into the active and inactive ports). Twenty-five minutes prior to the test, rats received either an injection of CNO (5 mg/kg i.p.) to activate the G_i_ DREADDs or vehicle (6% DMSO in 0.9% sterile saline) in an adjacent testing room. After the rats were placed in the testing chambers, one minute elapsed prior to illumination of the house light. The cue light in the active port also came on at the session start, and remained on for 20 seconds, as it did during the cocaine self-administration procedure. Pokes into the active port were recorded and resulted in presentation of the cue light for 5-sec, but no cocaine infusion, while pokes into the inactive port were recorded but without consequence. The test lasted for 2 h and occurred between 10:00 h and 17:00 h.

#### Cocaine-induced reinstatement test

Rats underwent 5 days of forced abstinence followed by extinction training 2 (8 sessions) prior to the cocaine-induced reinstatement test. As with the cue-induced reinstatement test, rats received the same treatment (i.e. CNO or vehicle) in an adjacent testing room 25-min prior to the test session. Before being put into the testing chamber, all rats received a 15 mg/kg (i.p.) injection of cocaine. This dose of cocaine was chosen as it has previously been shown to result in ST/GT differences in drug-seeking behavior during a test for cocaine-induced reinstatement [17]. The house light came on one minute after the session was initiated, but in contrast to the cue-induced reinstatement test the cue-light did not come on at any point during the test session. Pokes into the active and inactive port were recorded, but without consequence (i.e. no cue-light presentation when pokes were made into the active port). The session lasted for 2 h and occurred between 10:00 h and 17:00 h.

#### Histology

Fos protein expression was analyzed in a subset of rats to verify that the G_i_ DREADDs were in fact affecting neuronal activity in the PrL-PVT pathway. Rats received either CNO or vehicle (same as test day treatment) eighty minutes prior to perfusions, but did not undergo any additional tasks or experimentation. Thus, we were assessing the effects of DREADD activation (i.e. PrL-PVT pathway inhibition) on baseline neuronal activity (or that in response to an injection), as rats remained in the colony room after receiving treatment. This time point was chosen to capture peak Fos levels [1]. Co-localization of Fos positive cells and cells expressing the Gi DREADD in the PrL were manually quantified using ImageJ (National Institute of Health, Bethesda, MD) to determine selectivity of neuronal activation in response to CNO. To assess the effects of PrL-PVT inhibition on activity in the PVT, the density of Fos (cells per square mm) was determined using the ITCN (image-based tool for counting nuclei) plugin from ImageJ.

#### Tissue Processing

Once testing was complete, all rats were anesthetized with ketamine (90 mg/kg. i.p.) and xylazine (10 mg/kg, i.p.) and underwent transcardial perfusion. Rats were first perfused with 0.9% saline followed by 4% formaldehyde (pH= 4.7) at 4°C. After extraction, brains remained in formaldehyde for a 24 h period. Brains were then moved to graduated sucrose solutions (10%, 20% then 30% sucrose in phosphate buffer, pH= 7.4, 4°C) every 24 hours for 3 days for cryoprotection. Following cryoprotection, brains were encased in Tissue-Plus O.C.T. cryoprotectant (Fisher HealthCare, Houston, TX) and frozen using dry ice. They were then sliced at a thickness of 30 μm on a cryostat and underwent free-floating immunohistochemistry (IHC) for mCherry (protein fused to DREADD) to assess hM4D(G_i_) DREADD expression, and a subset of brains underwent an additional IHC for Fos protein expression.

Following IHC, brains were mounted and dehydrated in ethanol solution, washed in xylene and coverslipped with Permount (Fisher Scientific, Fair Lawns, NJ). Using a Leica DM1000 light microscope (Buffalo Grove, IL), DREADD expression in the PrL, aPVT and pPVT were qualitatively scored by two researchers who were blind to group assignments. DREADD expression scores ranged from 0 to 3 and were based on localization and strength of the expression. In order to be included in final analysis (scores of 1-3), DREADD expression had to be localized to the PrL (relative to bregma: AP 5-2.50), aPVT (relative to bregma: AP ^−^1.20-^−^2.50) and pPVT (relative to bregma: AP ^−^2.51-^−^3.50), and had to be easily visible under the microscope. A score of 1 indicated weak expression, whereas a score of 3 indicated strong expression. Rats with a score of zero (e.g. inaccurate injection, lack of expression in a region, etc.) were not included in final analyses. See Figure S1 for representation of viral spread within the PrL, aPVT and pPVT regions.

#### Immunohistochemistry (IHC)

##### DREADD expression for inclusion criteria

All procedures occurred at room temperature on a lab shaker on a low speed. Slices were washed in 0.1 M phosphate-buffered saline (PBS, pH= 7.3-7.4) five times (5-min each wash) and then blocked in 1% hydrogen peroxide in PBS solution for 10-min. Slices were then washed in PBS (4 washes) and blocked in 0.4% Triton-X and 2.5% Normal Donkey Serum (Jackson ImmunoResearch, West Grove, PA) diluted in PBS for one hour, and then incubated overnight in a PBS solution containing 0.4% Triton-X and 1% Normal Donkey Serum with rabbit anti-mCherry primary antibody (Abcam, Cambridge, MA) at a 1:30,000 dilution. The next day, slices were washed three times in PBS and incubated for one hour in a PBS solution containing biotinylated donkey anti-rabbit secondary antibody (Jackson ImmunoResearch, West Grove, PA) at a 1:500 dilution, 0.4% Triton-X, and 1% Normal Donkey Serum. After another series of three PBS washes, slices were incubated for one hour in ABC-elite (Vector Laboratories, Burlingame, CA) diluted at 1:1000 in PBS, and then underwent three PBS washes. In order to view DREADD expression, slices were then incubated in a 0.1M sodium phosphate-buffered solution containing 0.02% 3, 3’ diaminobenzidine tetrahyrochloride (DAB; Sigma Aldrich, St. Louis, MO) and 0.012% hydrogen peroxide. Incubation lasted for 8-min, and then slices were rinsed three times in PBS and stored at 4°C until being mounted. DAB staining creates a brown residue at the location of DREADD expression, allowing for subsequent visualization.

##### Fos protein expression

Fos protein expression in response to test day treatment (CNO or vehicle) was analyzed in a subset of rats (GT VEH: 10, GT CNO: 8, ST VEH: 3, ST CNO: 6) to verify that the G_i_ DREADDs were in fact affecting the activity in the PrL-PVT pathway. Eighty minutes prior to perfusions, rats were injected (i.p.) with either 5.0 mg/kg CNO or vehicle (same as test day treatment) and left in the colony room. This time period was chosen as Fos protein expression has been shown to peak 60-90-min after cellular activation [1]. Brains from rats with good DREADD expression underwent free-floating IHC for co-expression of Fos protein and DREADD expression. The IHC for Fos protein was performed first, and all procedures are the same as indicated above with the following exceptions: a goat anti-Fos primary antibody (Santa Cruz Biotechnology, Dallas, TX) diluted at 1:1000 and a biotinylated donkey anti-goat secondary antibody (Jackson ImmunoResearch, West Grove, PA) diluted at 1:500 were used. Additionally, nickel sulfate (0.08%) was added to the DAB solution so that Fos protein expression would have a black, and not brown residue, allowing for contrast between the two stains. Immediately following the IHC for Fos protein, the IHC for DREADD expression was completed as described above.

To assess the effects of PrL-PVT inhibition on activity in the PVT, Fos protein expression was quantified in the PVT. One PVT section in the anterior (relative to Bregma: ^−^2.00), middle (relative to Bregma: AP ^−^2.50), and posterior (relative to Bregma: AP ^−^3.00) subregions was analyzed using ImageJ (National Institute of Health, Bethesda, MD). The number of Fos positive cells and density of expression (cells per square mm) were determined using the ITCN (image-based tool for counting nuclei) plugin from ImageJ.

##### DREADD expression for representative images

A select number of rats also underwent an IHC to fluorescently label DREADD expression for image presentation in this manuscript (see Figure 1). Slices were washed in PBS three times (5-min each wash) and then blocked in 0.4% Triton-X and 2.5% Normal Donkey Serum (Jackson ImmunoResearch, West Grove, PA) diluted in PBS for one hour. An overnight incubation then occurred in a PBS solution containing 0.4% Triton-X and 1% Normal Donkey Serum with rabbit anti-mCherry primary antibody (Abcam, Cambridge, MA) at a 1:1000 dilution. The following day, slices underwent three PBS washes, a two-hour incubation in a PBS solution containing 0.4% Triton-X and 1% Normal Donkey Serum with biotinylated donkey anti-rabbit secondary antibody (Jackson ImmunoResearch, West Grove, PA) at a 1:500 dilution, followed by three more PBS washes. The remainder of the IHC was done under minimal light conditions. Slices were incubated for one hour with Alexa FluorTM 594-conjugated streptavidin (Thermo Fisher Scientific, Waltham, MA) diluted 1:1000 in PBS with 0.4% Triton-X, then 3 more PBS washes. Next, slices were incubated for 20-min in DAPI diluted 1:2000 in PBS and then mounted onto slides and coverslipped with ProLong Gold Antifade Mountant (Thermo Fisher Scientific, Waltham, MA). Images were taken using an Olympus FV3000 Confocal Laser Scanning Microscope.

#### Video analysis: Cocaine-primed reinstatement test

To assess if the induction of stereotyped behaviors contributed to the variance of behavior observed during the cocaine-induced reinstatement test, both stereotyped and non-stereotyped behaviors for each rat was quantified. Quantified behavior was selected due its high presence shown by several rats when videos were randomly sampled for screening. Stereotyped behaviors include: circling, head movements but no port entry, head movements and port entry. Non-stereotyped behaviors include: immobility, orientation toward the active port without movement toward it, approach to the active port without port entry. A more detailed description of the criteria for each behavior can be found in Table S2. Behavior was analyzed for 30-sec every ten minutes for the first hour of the reinstatement test, as this is when peak drug-seeking behavior occurred. The percentage of time (min) rats spent exhibiting each behavior across the six bins was quantified.

#### Statistical analyses

To assess if there was a relationship present between rate of extinction (i.e. decrease in active nose pokes) and the reinstatement tests, a quadratic regression model was fit to each rat’s extinction training 1 and 2 curve. The quadratic term was then regressed onto the number of nose pokes into the active port during the reinstatement test that followed each extinction training. This analysis allows us to account for differences in drug-seeking behavior during the reinstatement tests as a function of rate of extinction. SPSS version 22 was used for these analyses and statistical significance was set at p<0.05.

All significant results for the tests for cue-induced and cocaine-primed reinstatement were subjected to a power analysis and exhibited adequate statistical power (1-β>0.55).

#### Histological analyses: PrL

Group differences in the number of Fos positive cells, DREADD positive cells, and cells expressing both markers were analyzed using an ANOVA. The percentage of cells co-expressing DREADD and Fos relative to all cells expressing DREADD was analyzed using an ANOVA.

#### Histological analyses: PVT

The average density of Fos positive cells collapsed across the rostral-caudal axis of the PVT were assessed using an ANOVA. Rats in the vehicle group, regardless of phenotype, were collapsed into a single group as they did not statistically differ from one another. An unpaired t-test was used to assess differences between CNO-treated STs and the vehicle group, and CNO-treated GTs and the vehicle group. To further assess if PrL-PVT inhibition affected Fos density in the PVT of STs and GTs, the percent of Fos relative to all vehicle control rats (i.e. “baseline”) was calculated for each phenotype. A single sample t-test with the hypothesized value set to 100% (i.e. baseline) was then performed. To determine if PrL-PVT inhibition differentially affected PVT Fos density relative to baseline in STs versus GTs, an unpaired t-test was performed.

### Results

#### Subjects

As indicated in the primary text, only rats that successfully completed all training and exhibited accurate DREADD expression were included in final analysis. Figure S1 illustrates the detected spread of DREADD expression for those rats that were included. Though expression in the aPVT appears more diffuse, a majority of the expression is within the boundaries of the aPVT. Additionally, expression in the PrL and pPVT are heavily localized, suggesting off target expression into the aPVT is likely unavoidable.

Seven rats died during catheterization or stereotaxic surgery, prior to training. Three rats were eliminated from the study following PavCA training as they did not acquire the task (i.e. did not approach the lever-CS or food magazine during training). One rat was sacrificed at the conclusion of food self-administration training due to health complications. Additional rats were removed from the study if they did not meet the IC during self-administration training (n= 51 (ST: 4/36; GT: 29/96) or due to loss of catheter patency, n= 1 (ST)).

### Behavioral results: ST/GT model

#### More STs acquire cocaine self-administration than GTs

More GTs (30%) were excluded from the study due to not meeting IC requirements compared to STs (11%). This suggests that while behavior did not differ between rats that did acquire cocaine self-administration, STs are more likely to acquire self-administration compared to GTs. While this is true in the current study, prior studies using the same self-administration paradigm did not observe this trend [2–4].

#### All rats exhibited cue-induced reinstatement and inhibition of the PrL-PVT pathway selectively attenuates drug-seeking behavior in STs

To compare responding during the cue-induced reinstatement test to that during the last extinction session, we examined the effects of Phenotype (ST or GT) and Treatment (VEH or CNO) on the number of NPs in the active port minus those in the inactive port (i.e. active – inactive nose pokes). For this dependent variable, all rats showed an increase in drug-seeking behavior during the test for cue-induced reinstatement compared to the last extinction training session (effect of Session, F_1,40_=33.37, p<0.001; Figure 4b). There was a significant effect of Phenotype (F_1,40_=4.59, p=0.04) and Treatment (F_1,40_=7.51, p=0.009). A significant Treatment x Session interaction (F_1,40_=5.52, p=0.02) and Treatment x Phenotype interaction (F_1,40_=6.48, p=0.01) were also present, indicating that responding was differentially affected in a treatment-dependent manner between the two sessions and two phenotypes (Figure 4b). Using this metric (active-inactive nose pokes) we also assessed the relationship between cue-induced reinstatement and PavCA index for each of the experimental groups, and found no significant correlations (ST VEH, r^2^=0.75, p= 0.06; ST CNO, r^2^=0.13, p= 0.38; GT VEH, r^2^<0.001, p=0.99; GT CNO, r^2^=0.05, p= 0.40). As sample size likely impacted these results, the trend level of significance reached in the ST VEH group suggests that the degree to which one sign-tracks may predict cue-induced reinstatement of drug-seeking behavior, and that manipulation of the PrL-PVT pathway perturbs this relationship.

#### All rats exhibited cocaine-primed reinstatement

Drug-seeking behavior (active-inactive nose pokes) between the last extinction session and the test for cocaine-induced reinstatement was analyzed as a means to directly compare responses between the two sessions. All rats showed greater drug-seeking behavior during the test for cocaine-induced reinstatement compared to the last extinction training session (effect of Session, F_1,40_=34.48, p<0.0001; Figure 4i). There was not a significant effect of Treatment (effect of Treatment, F_1,40_=0.06, p=0.81), Phenotype (effect of Phenotype, F_1,40_=0.01, p=0.93), nor any significant interactions present for this dependent variable during the test for cocaine-induced reinstatement. There were no significant correlations between nose pokes made into the active-inactive port during the tests for cue-induced reinstatement and cocaine-primed reinstatement (ST VEH, r^2^=0.12, p= 0.57; ST CNO, r^2^=0.03, p= 0.72; GT VEH, r^2^=0.23, p=0.07; GT CNO, r^2^<0.001, p= 0.96); nor was there a significant correlation between PavCA index score and cocaine-primed reinstatement (active-inactive nose pokes) in any of the groups (ST VEH, r^2^=0.51, p= 0.11; ST CNO, r^2^=0.02, p= 0.76; GT VEH, r^2^=0.11, p=0.23; GT CNO, r^2^=0.10, p= 0.26).

#### STs and GTs do not differ in cocaine-induced stereotyped behaviors during cocaine primed-reinstatement

Stereotyped behaviors during cocaine-primed reinstatement were assessed in a subset of rats (ST VEH, n=4; ST CNO, n=3; GT VEH, n=10; GT CNO, n=9). Rats spent more time exhibiting “non-stereotyped behaviors” (e.g. active port orientation/ approach, immobility) compared to “stereotyped behaviors” (e.g. head movements, circling) (effect of Category, F_1,22_=8.13, p<0.001; Figure S2). On average, rats spent ~5-10% percent of the time exhibiting stereotyped behaviors. STs and GTs did not differ on this measure (effect of Phenotype, F_1,22_<0.0001, p=0.99) and treatment did not affect cocaine-induced stereotypy (effect of Treatment, F_1,22_=0.04, p=0.84; Figure S2). These data suggest that while some animals exhibited stereotypy, this drug-induced response likely did not interfere with the observed differences (or lack thereof) during the cocaine-induced reinstatement test. However, further investigation is warranted given the small sample size.

#### Rate of extinction does not predict reinstatement of drug-seeking behavior in STs or GTs

There was not a significant relationship present between the rate of extinction (extinction 1; i.e., the change in responding in the active nose port across sessions) and the number of nose pokes made into the active port during the test for cue-induced reinstatement (ST VEH, r^2^=0.20, p=0.45; ST CNO, r^2^=0.12, p=0.41; GT VEH, r^2^=0.01, p=0.79; GT CNO, r^2^=0.05, p=0.45). This suggests that the rate by which rats decreased nose pokes made into the active port during extinction training did not correlate with cue-induced reinstatement of drug-seeking behavior. There was also no relationship present between the rate of extinction (extinction 2) and the number of nose pokes made into the active port during cocaine-primed reinstatement (ST VEH, r^2^=0.41, p=0.17; ST CNO, r^2^=0.03, p=0.67; GT VEH, r^2^=0.02, p=0.67; GT CNO, r^2^=0.20, p=0.10), suggesting that variation in drug-seeking behavior seen during this test is not due to differences in extinction rate.

#### CNO in the absence of DREADD does not affect reinstatement of drug-seeking behavior

To control for potential effects of CNO on behavior, a separate cohort of rats (ST, n= 10; GT, n=15) underwent all experimental procedures as previously described in this manuscript, with the exception of undergoing stereotaxic surgery for DREADD virus expressions. During the reinstatement tests, rats in this group were administered either CNO or vehicle. All rats exhibited reinstatement of drug-seeking behavior relative to the last extinction training session during the test for cue-induced reinstatement (effect of Session, F_1,21_=15.97, p=0.001; Figure S3a) and cocaine-primed reinstatement (effect of Session, F_1,19_=58.81, p<0.0001; Figure S3b). Furthermore, rats made more pokes into the active port compared to the inactive port during both reinstatement tests (Cue: effect of Port, F_1,42_=15.17, p<0.001, Figure S3a; Prime: effect of Port, F_1,38_=42.26, p<0.0001, Figure S3b). Unlike prior studies, STs and GTs did not differ in either cue-induced (F_1,42_=0.003, p=0.95) [3, 5] or cocaine-primed (F_1,42_=0.003, p=0.95) [4] reinstatement of drug-seeking behavior. Importantly, administration of CNO prior to the test session in the absence of DREADDs did not affect drug-seeking behavior (Cue: effect of Treatment, F_1,42_=0.53, p=0.47, Figure S3a; Cocaine prime: effect of Treatment, F_1,38_=0.07, p=.80, Figure S3b).

### Behavioral results: High-responder (HR)/ Low-responder (LR) model

#### All rats exhibited cue-induced reinstatement

Compared to the last extinction training sessions, all rats showed greater drug-seeking behavior during the cue-induced reinstatement test compared to the last extinction training session when active-inactive nose pokes was used as the dependent variable (effect of Session, F_1,40_=31.89, p<0.0001; Figure 6b). There was a significant effect of Phenotype (F_1,40_=5.30, p=0.03) with HRs showing greater drug-seeking behavior compared to LRs (Figure 6b), but no significant effect of Treatment (F_1,40_=3.52, p=0.07) or any interactions. Nose pokes made into the active-inactive port during the reinstatement test did not significantly correlate with locomotor score for rats in any of the groups (HR VEH, r^2^=0.46, p=0.09; HR CNO, r^2^=0.26, p= 0.11; LR VEH, r^2^=0.001, p=0.92; LR CNO, r^2^=0.13 p= 0.26). Therefore, it appears locomotor activity in a novel environment does not predict levels of cue-induced reinstatement of drug-seeking behavior in HRs or LRs.

#### All rats exhibited cocaine-primed reinstatement

All rats exhibited greater drug-seeking behavior during the cocaine-prime reinstatement test compared to the last extinction training session when active-inactive nose pokes was used as the dependent variable (effect of Session, F_1,40_=39.78, p<0.0001; Figure 6i). There was not a significant effect of phenotype (effect of Phenotype, F_1,40_=0.25, p=0.62), treatment (effect of Treatment, F_1,40_=0.01, p=0.94), nor any significant interactions for this measure during the cocaine-primed reinstatement test. Furthermore, cumulative locomotor movements made during the locomotor test did not correlate with drug-seeking behavior (active-inactive nose pokes) for any of the groups (HR VEH, r^2^=0.01, p= 0.83; HR CNO, r^2^=0.09, p= 0.38.; LR VEH, r^2^=0.18, p= 0.15; LR CNO, r^2^=0.004, p= 0.85).

#### HRs and LRs do not differ in cocaine-induced stereotyped behaviors during cocaine primed-reinstatement

Stereotyped behaviors during cocaine-primed reinstatement were assessed in a subset of rats (HR VEH, n=5; HR CNO, n=8; LR VEH, n=9; LR CNO, n=4). Rats spent less time exhibiting “stereotyped behaviors” (~5-15% of time) compared to “non-stereotyped behaviors” (effect of Category, F_1,22_=9.33, p<0.01; Figure S4). HRs and LRs did not differ on this measure (effect of Phenotype, F_1,22_=0.22, p=0.65) and treatment did not affect stereotyped behavior (effect of Treatment, F_1,22_=0.29, p=0.59; Figure S4). Thus, in parallel to what was observed with the ST/GT model, it appears some animals exhibited stereotypy, but this likely did not interfere with cocaine-induced reinstatement measures. However, further investigation is warranted given the small sample size.

#### Rate of extinction does not predict reinstatement of drug-seeking behavior in HRs or LRs

There was not a significant relationship between the rate of extinction (extinction 1) and cue-induced drug-seeking behavior (HR VEH, r^2^=0.04, p=0.68; HR CNO, r^2^=0.02, p=0.68; LR VEH, r^2^<0.001, p=0.99; LR CNO, r^2^=0.13, p=0.26), or extinction rate (extinction 2) and cocaine-primed drug-seeking behavior (HR VEH, r^2^=0.11, p=0.41; HR CNO, r^2^=0.08, p=0.41; LR VEH, r^2^=0.06, p=0.43; LR CNO, r^2^=0.15, p=0.22) for any of the groups. Thus, the variation in drug-seeking behavior does not appear to be due to differences in extinction rate.

### Histology results

#### PrL Fos and DREADD expression

There were no significant effects of treatment, phenotype or interactions for the number of cells expressing DREADD (Treatment, F_1,22_=1.37, p=0.26; Phenotype: F_1,22_=3.25, p=0.09), Fos (Treatment: F_1,22_=0.08, p=0.78; Phenotype: F_1,22_=1.57, p=0.22), or co-expressing both Fos and DREADD (Treatment: F_1,22_=0.54, p=0.47; Phenotype: F_1,22_=2.65, p=0.19) in the PrL. As there was no effect of Phenotype in any analysis, STs and GTs were collapsed together for further analysis. Relative to vehicle, CNO administration resulted in a significant increase in the percent of cells co-expressing both Fos and DREADD relative to all cells expressing DREADD (F_1,22_=5.8, p=0.02; Figure S5a). Thus, CNO administration increased neuronal activity in DREADD-expressing cells in the PrL. Though this appears counterintuitive to what is to be expected, it should be emphasized that this was merely a test of CNO-induced changes in neuronal activity, as rats were left undisturbed in their colony room prior to sacrifice and did not undergo any task. Thus, it is possible that either under basal conditions, or during normal ambulatory behavior, the PrL-PVT pathway is inhibited and thus chemogenetic inhibition of the pathway results in a disinhibition and increase in neuronal activity. However, further investigation is needed to assess this hypothesis.

#### PVT Fos expression

Density of Fos protein expression in the PVT was analyzed across PVT sub-regions (anterior, middle and posterior), but there was not a significant effect of Region (F_2,40_=1.31, p=0.28) nor a significant Phenotype x Treatment x Region interaction (F_2,40_=0.37, p=0.70). Therefore, all subsequent analyses focused on the average density of Fos across the rostral-caudal axis of the PVT, for which there was a significant Phenotype x Treatment interaction (F_1,23_= 4.29, p<0.05; Figure S5b). Post-hoc analyses showed that inhibition of the PrL-PVT circuit differentially affected average Fos density in STs relative to GTs (p=0.002, Cohen’s d=2.05), such that average Fos density decreased in GTs treated with CNO relative to STs treated with CNO (Figure S5b). However, when compared to their respective vehicle-treated controls, PrL-PVT inhibition did not affect average Fos density in STs (p=0.19, Cohen’s d=0.92) or GTs (p=0.10, Cohen’s d=0.87). Because there was a not a significant difference in average Fos density between the vehicle-treated groups (p=0.49, Cohen’s d=0.50), additional analyses were conducted with a single vehicle group collapsed across phenotypes.

When the percent of average Fos density relative to baseline (i.e. collapsed vehicle control group) was analyzed, PrL-PVT inhibition resulted in greater PVT Fos density relative to baseline in STs compared to GTs (t(12)=−4.10, p<0.002; Cohen’s d=2.05; Figure S5c). These findings are consistent with the results reported above using the raw data. Relative to baseline (i.e. 100%), inhibition of the PrL-PVT did not significantly affect Fos in the PVT in STs (t(5)=1.36, p=0.23), but did decrease Fos in the PVT in GTs (t(7)=^−^9.01, p<0.001, Figure S5c). However, there was no correlation between average Fos density relative to baseline and active lever presses during cue-induced reinstatement in either phenotype (GT: p=0.68, r^2^=0.03; ST: p=0.34, r^2^=0.23). These results support the notion that the PrL-PVT pathway differentially mediates downstream activity STs and GTs. However, as rats did not undergo a challenge to evoke neuronal activity in the current study, further investigation is warranted.

---

**Table S1.** Acquisition of sign- and goal-tracking behavior across PavCA training. STs exhibited more lever-directed behaviors over the course of PavCA training compared to GTs. Accordingly, GTs showed more magazine-directed behaviors compared to STs.

| *Lever-directed behaviors (Sign-tracking)* |  |  |  |  |  |  |  |  |  |
| --- | --- | --- | --- | --- | --- | --- | --- | --- | --- |
|  | Probability lever |  |  | Lever contacts |  |  | Latency lever |  |  |
|  | DF | F | P | DF | F | P | DF | F | P |
| *Session* | 4,42 | 25.25 | **<0.0001** | 4,42 | 18.22 | **<0.0001** | 4,42 | 35.16 | **<0.0001** |
| *Phenotype* | 1,42 | 183.67 | **<0.0001** | 1,42 | 85.04 | **<0.0001** | 1,42 | 127.60 | **<0.0001** |
| *Treatment* | 1,42 | 0.48 | 0.49 | 1,42 | 1.28 | 0.26 | 1,42 | 0.67 | 0.42 |
| *Session\*Phenotype* | 4,42 | 28.68 | **<0.0001** | 4,42 | 17.89 | **<0.0001** | 4,42 | 34.62 | **<0.0001** |

| *Magazine-directed behaviors (Goal-tracking)* |  |  |  |  |  |  |  |  |  |
| --- | --- | --- | --- | --- | --- | --- | --- | --- | --- |
|  | Probability magazine |  |  | Magazine contacts |  |  | Latency magazine |  |  |
|  | DF | F | P | DF | F | P | DF | F | P |
| *Session* | 4,42 | 4.02 | **0.005** | 4,42 | 4.52 | **0.002** | 4,42 | 4.59 | **0.002** |
| *Phenotype* | 1,42 | 55.81 | **<0.0001** | 1,42 | 76.83 | **<0.0001** | 1,42 | 61.62 | **<0.0001** |
| *Treatment* | 1,42 | 0.87 | 0.36 | 1,42 | 0.13 | 0.72 | 1,42 | 0.52 | 0.48 |
| *Session\*Phenotype* | 4,42 | 11.25 | **<0.0001** | 4,42 | 13.66 | **<0.0001** | 4,42 | 11.61 | **<0.0001** |

**Table S2.** Table of behaviors quantified by a blind observer during video analysis of the test for cocaine-primed reinstatement. Behaviors were scored in 30-sec bins every 10-min for the first 60-min of the session. Time spent in each behavior was recorded per bin.

| Behavior | Definition |
| --- | --- |
| *Stereotyped behaviors* |  |
| Circling (i.e. "spinning") | Four consecutive 90° turns occurring within close succession of one another |
| Head movements | Lateral head movements made when the back paws remain on the flooring and in the same position for a minimum of 2-sec |
| *Non-stereotyped behaviors* |  |
| Active port orientation | Oriented toward the active port, but not advancing toward the port |
| Active port approach | Approached the active port, but made no pokes into the port |
| Immobility | Time during which all 4 paws remain on the flooring in the same position for a minimum of 2-sec, which no other behaviors co-occurring |

**Figure S1.**
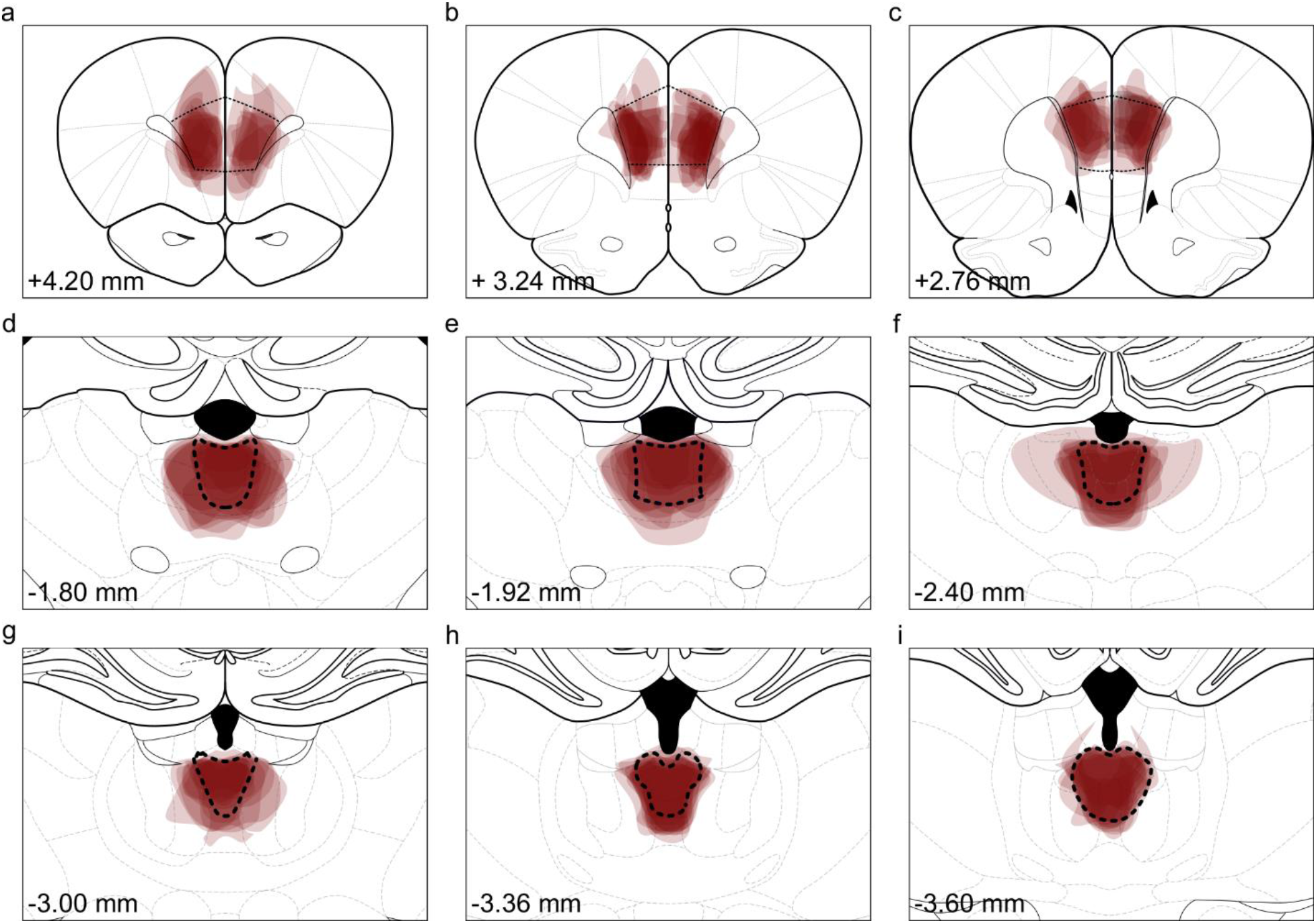
DREADD viral expression patterns in the PrL, aPVT and pPVT. Schematics showing DREADD viral vector spread within the **a-c)** PrL (relative to bregma: AP 4.20-2.76), **d-f)** aPVT (relative to bregma: AP ^−^1.80-^−^2.40) and **g-i)** pPVT (relative to bregma: AP ^−^3.00-^−^3.60). The intensity of red coloration on the schematics represents the number of rats exhibiting that viral spread pattern. The darker the red, the greater the number of rats. Rats included in final analyses were assigned a score of 1-3, with 3 being those with the best expression and localization. Those without sufficient expression and localization were excluded from final analyses.

**Figure S2.**
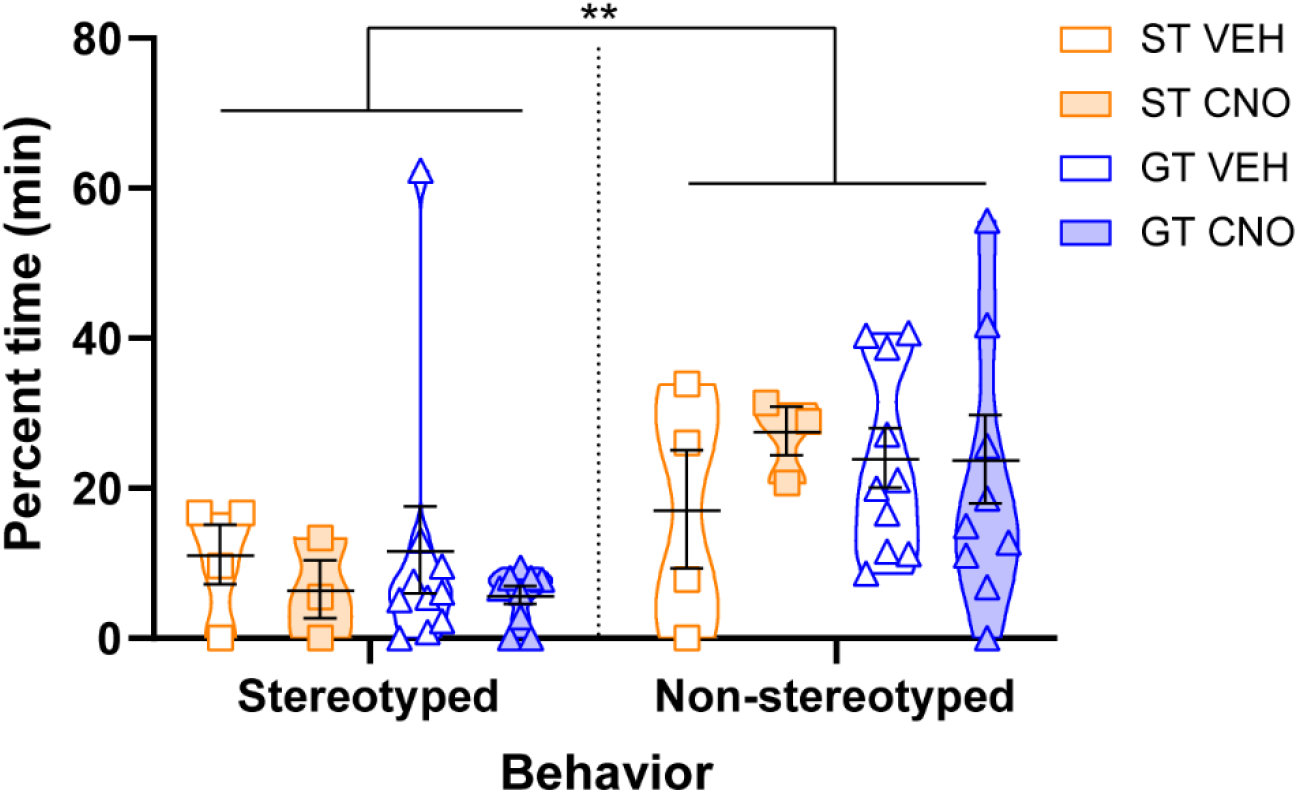
Stereotyped behavior and non-stereotyped behavior during cocaine-primed reinstatement. Data points illustrate the percent of time each rat spent in stereotyped or non-stereotyped behaviors during the first hour of the test for cocaine-primed reinstatement. Black bars within violin plots represent the mean ± SEM for percent time exhibiting stereotyped behavior. All rats exhibited more non-stereotyped than stereotyped behavior (p<0.001), and behavior did not differ based on phenotype (p=0.99) or treatment (p=0.84). (ST VEH, n=4; ST CNO, n=3; GT VEH, n=10; GT CNO, n=9) **p<0.01

**Figure S3.**
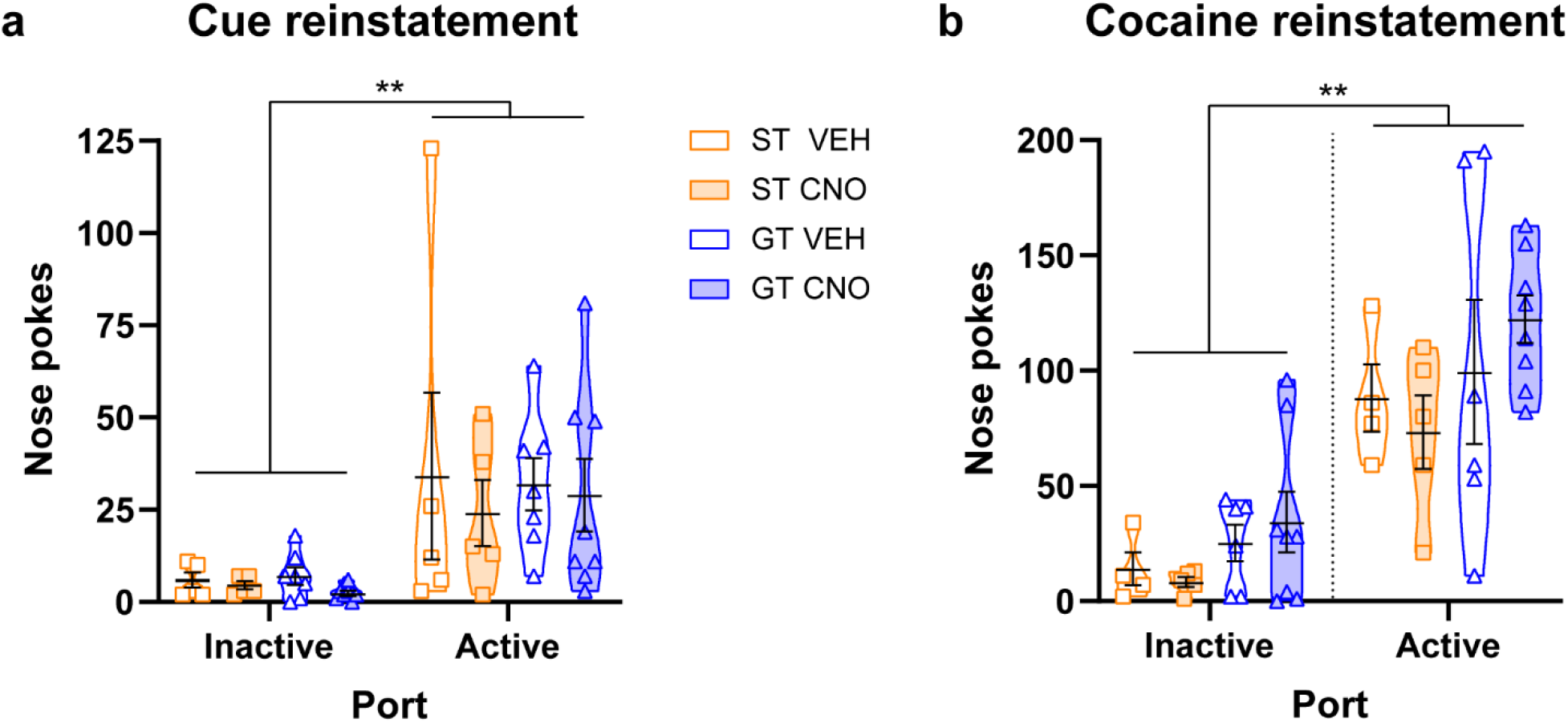
CNO administration in the absence of DREADDs did not affect drug-seeking behavior during reinstatement. Individual data points represent mean number of nose pokes into the active and inactive port with black bars within violin plots representing the mean ± SEM during **a)** a test for cue-induced reinstatement and **b)** a test for cocaine-primed reinstatement. CNO administration in the absence of DREADDs did not affect drug-seeking behavior during either reinstatement test in STs or GTs (Cue: F_1,42_=0.53, p=0.47; Prime: F_1,38_=0.07, p=.80). (ST VEH, n=5 (n=4 for cocaine reinstatement); ST CNO, n=5; GT VEH, n=7 (n=6 for cocaine reinstatement); GT CNO, n=8)

**Figure S4.**
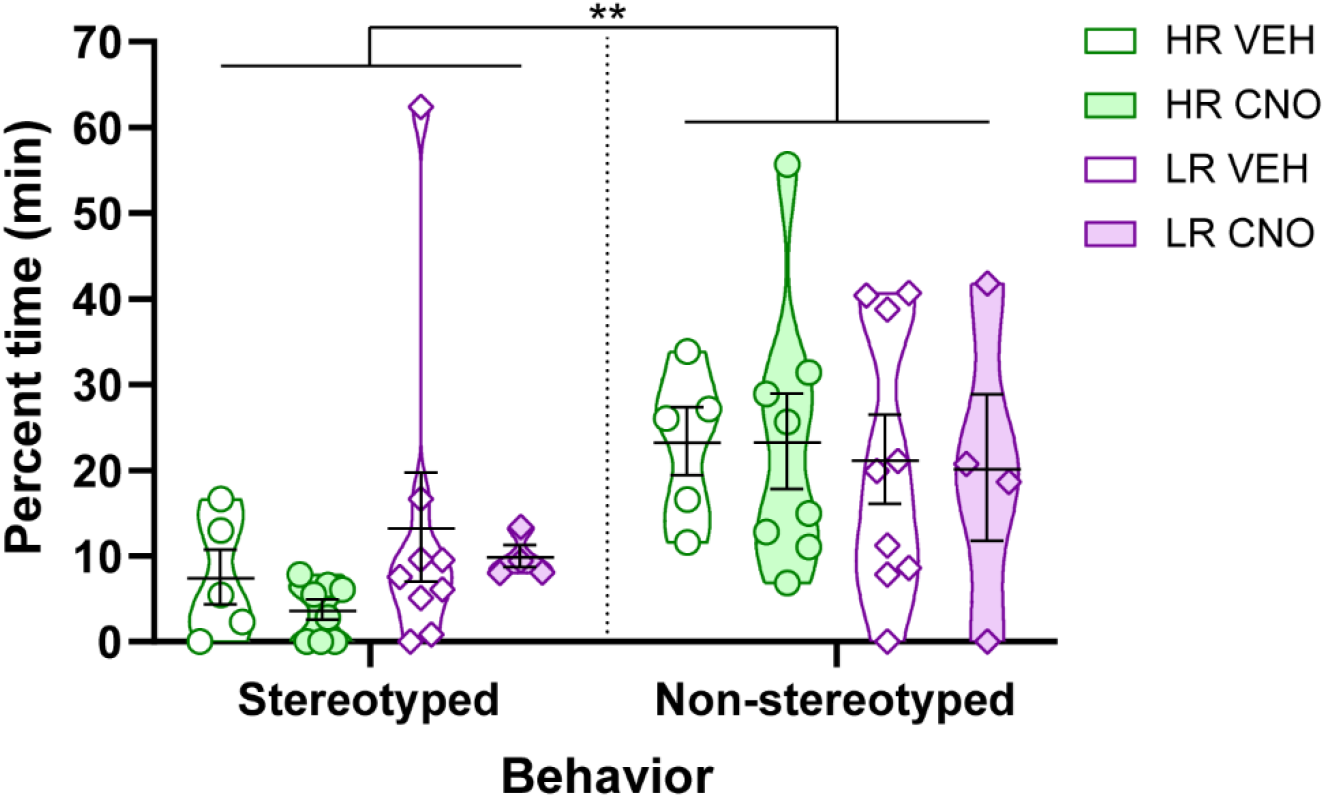
Cocaine-induced stereotyped behavior during cocaine-primed reinstatement. Data points illustrate the percent of time each rat spent in stereotyped or non-stereotyped behaviors during the first hour of the test for cocaine-primed reinstatement. Black bars within violin plots represent the mean ± SEM for percent time exhibiting stereotyped behavior. Behavior did not differ based on phenotype (p=0.59) or treatment (p=0.76). (HR VEH, n=5; HR CNO, n=8; LR VEH, n=9; LR CNO, n=4)

**Figure S5.**
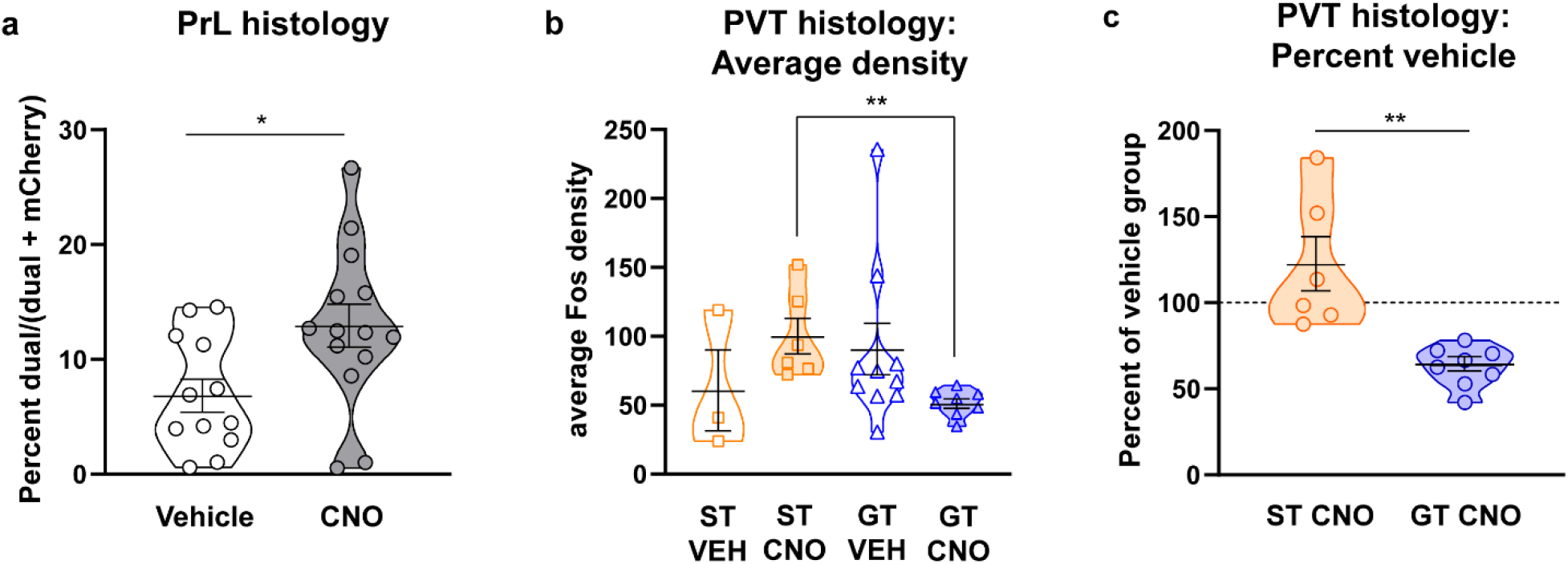
Histology of cells in the PrL and PVT of STs and GTs. **a)** Individual data points shown for the percent of cells across the PrL (anterior, middle and posterior) co-expressing both Fos and mCherry (DREADD tag) in relation to all cells expressing mCherry. Mean ± SEM shown as black bars within violin plots. CNO administration increases this percentage compared to vehicle treated rats (p=0.02) (VEH group, n=12 (ST, n=2; GT, n=10); CNO group, n=14 (ST, n=6; GT, n=8)). **b)** Individual data points represent the average Fos density in the PVT (anterior, middle and posterior) and black bars within the violin plots represent the mean ± SEM. Inhibition of the PrL-PVT pathway results in a decrease in average Fos density in GTs relative to STs (p=0.002). However, inhibition of the PrL-PVT pathway does not affect average Fos density in either phenotype with respect to their vehicle-treated controls (ST: p= 0.19; GT, p= 0.10), and rats in the vehicle groups did not differ from one another (ST VEH relative to GT VEH, p=0.49). **c)** Individual data points shown for the percent Fos density throughout the entire PVT (anterior, middle and posterior). Mean ± SEM shown as black bars within violin plots. Pooled vehicle-treated group (i.e. baseline) indicated by dashed line at 100%. Inhibition of the PrL-PVT pathway results in a decrease in Fos density in GTs relative to vehicle-treated rats (p<0.001), and no effect in STs relative to vehicle-treated rats (p=0.23). Inhibition of the PrL-PVT pathway differentially affected PVT Fos density (% vehicle) in STs relative to GTs (p=0.002; **p<0.01). (ST VEH, n=3; ST CNO, n=6; GT VEH, n=10; GT CNO, n=8) **p<0.01

